# redPATH: Reconstructing the Pseudo Development Time of Cell Lineages in Single-Cell RNA-Seq Data and Applications in Cancer

**DOI:** 10.1101/2020.03.05.977686

**Authors:** Kaikun Xie, Zehua Liu, Ning Chen, Ting Chen

## Abstract

Recent advancement of single-cell RNA-seq technology facilitates the study of cell lineages in developmental processes as well as cancer. In this manuscript, we developed a computational method, called redPATH, to reconstruct the pseudo developmental time of cell lineages using a consensus asymmetric Hamiltonian path algorithm. Besides, we implemented a novel approach to visualize the trajectory development of cells and visualization methods to provide biological insights. We validated the performance of redPATH by segmenting different stages of cell development on multiple neural stem cell and cancerous datasets, as well as other single-cell transcriptome data. In particular, we identified a subpopulation of malignant glioma cells, which are stem cell-like. These cells express known proliferative markers such as *GFAP* (also identified *ATP1A2*, *IGFBPL1*, *ALDOC*) and remain silenced in quiescent markers such as *ID3*. Furthermore, *MCL1* is identified as a significant gene that regulates cell apoptosis, and *CSF1R* confirms previous studies for re-programming macrophages to control tumor growth. In conclusion, redPATH is a comprehensive tool for analyzing single-cell RNA-Seq datasets along a pseudo developmental time. The software is available via http://github.com/tinglab/redPATH.

## Introduction

Developmental research at a single cell level has been supported by flow cytometry and imaging methods over the past few decades. Three fundamental questions of interest include how individual cells develop into different cell types and tissues, how these cells function, and the underlying mechanism in gene regulations. Such cell development processes have yet remained significantly obscure [1]. Recent advances in single-cell RNA sequencing (scRNA-seq) technology [2] have enabled us to characterize the whole transcriptome of individual cells, thus allowing us to study the subtle difference in heterogeneous cell populations. For example, single-cell analysis in tumors, immunology, neurology, and hematopoiesis have led to new and profound biological findings [3–8].

Specifically, for glioblastoma (GBM), single-cell analysis reveals the functionality of tumor microenvironment in GBM. The relationships among microglia/macrophages, malignant cells, oligodendrocytes, and T cells have been uncovered, confirming previous biological conclusions [3–5]. Glioma associated microglia/macrophages (GAM) were known to regulate tumor growth, adversely changing its functionality under normal conditions [9–13]. Recent research [13] targeted GAM for re-activation of its antitumor inflammatory immune response to suppress tumor growth. Previously, potential markers (such as *CSF1R*) have also been identified for reprogramming of GAM; however, it acquired resistance over time and resumed vigorous tumor growth [14,15].

Many algorithms have been developed to study cell development processes, including cell differentiation and cell proliferation, by inferring a pseudo-time trajectory at the single-cell level for both snapshot data as well as multiple time-point data. A recent review [16] compared multiple state-of-the-art methods for developmental trajectory inference including Monocle2 [17], TSCAN [18], SCORPIUS [19], and for cell cycle processes such as reCAT [20]. Popular trajectory tools also include Seurat, DPT, Wishbone and numerous others [21–25]. Both Monocle2 and TSCAN assume a free branching structure of cell fate development whereas SCORPIUS assumes a linear development. Most of the existing methods would have two main steps, linear or non-linear dimensionality reduction followed by trajectory inference.

Monocle2 [17] uses an unsupervised feature selection called ‘dpFeature’, where it selects the genes that are differentially expressed among unsupervised clusters of cells. Then a principal graph is learned via a reverse graph embedding (RGE) algorithm, ‘DDRTree’, where it reflects the structure of the graph in much lower-dimensional space. The pseudo-time is then inferred by calculating a minimum spanning tree (MST) on the distance of the projection points to the line segment on the principal graph.

TSCAN [18] takes into consideration the dropout event. The raw gene expression is first processed by gene clustering to gain an average gene expression. Since many of the gene clusters are highly correlated, TSCAN reduces the dimensionality using principal component analysis (PCA). Then MST is applied to cell cluster centroids, which are inferred from the reduced space to form a trajectory. Finally, each cell is projected onto the MST trajectory to obtain the pseudo-time.

SCORPIUS [19] is a fully unsupervised trajectory inference method. First, it calculates the Spearman’s rank correlation between cells and defines an outlier metric for each cell. Then multi-dimensional scaling (MDS) is applied to the correlation distance matrix to learn a low dimensional representation of each cell. An initial principal curve is then calculated as the shortest path between the k-means (k set to 4) cluster centers of the cells. The principal curve is then learned iteratively by projecting the cells onto the curve.

Traditional trajectory inference analysis would remove cell cycle effects through the removal of cycling genes. However, the cell cycle process and cell differentiation process seem to be coupled according to recent research, especially in the development of neural stem cells (NSCs) [26]. Within the sub-ventricular zone (SVZ), it is estimated that 80% of adult neural stem cells undergo symmetric differentiation, and 20% undergo symmetric proliferation with little evidence of asymmetric divisions. To date, only one computational method, CycleX [27], attempts to decipher such a relationship between the two developmental processes.

In this work, redPATH successfully recovers the pseudo time of the differentiation process and also discovered some unique genes along with cell development (**Figure 1**). The performance and stability of redPATH are validated and compared with multiple state-of-art methods, showing its consistency in explaining marker gene expression changes. Here, we first implemented a consensus Hamiltonian path (cHMT) algorithm to reconstruct the pseudo-time of a linear differentiation process. We propose to model the differentiation process between cells using an asymmetric measure (Kullback-Leibler distance). The linear development assumption has importance in studies of more differentiated lineages at later stages of development.

**Figure 1.**
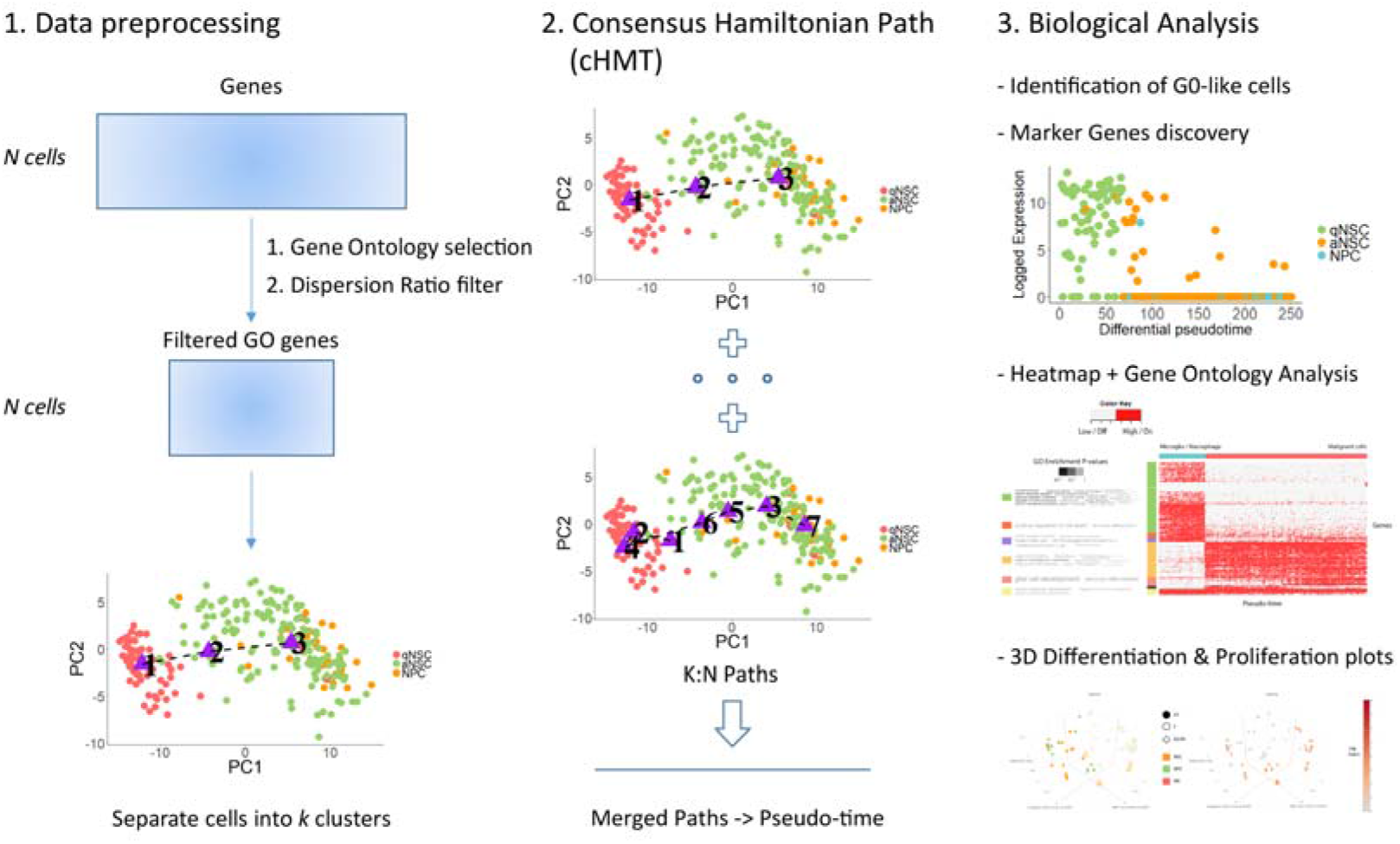
Overview of redPATH. Pipeline of the redPATH algorithm and analysis. Parts 1-2 provides the schematic illustration of the algorithm, comprising of data preprocessing steps and trajectory inference. The rightmost panel lists the main biological analysis functions included in redPATH.

Furthermore, the linear structure allows us to study the relationship between differentiation and proliferation more clearly. Additionally, we developed an approach to decipher and combine multiple Hamiltonian path solutions into a transition matrix to visualize the trajectory (linear or branched) developmental trend. Subsequent analysis and visualizations are implemented to provide biological insights into the developmental processes. redPATH is incorporated with reCAT in an attempt to visualize the relationship between differentiation and the cell cycle within the cell development of neural cells. Finally, glioma datasets are analyzed with a new perspective, uncovering a subpopulation within malignant cancerous cells.

## Materials and methods

### Overview of redPATH

As shown in **Figure 1**, redPATH consists of three main steps, namely data pre-processing, pseudo time inference, and biological analysis. There are two main challenges in the pseudo time inference problem in single-cell transcriptome data. The curse of dimensionality and the high level of noise together can severely affect the performance of pseudo time inference.

There are two main assumptions for redPATH. First, we assume the higher similarity between cells within the same cell type or the same developmental stage than those between different states. Second, although a linear developmental trend is assumed, multiple Hamiltonian path solutions are utilized to detect both linear and branching trajectories.

### Data pre-processing

The pre-processing step includes standard normalization procedures using existing methods such as edgeR and DEseq2 if the gene expression matrix is not yet normalized [28–31]. We take the log2 expression value for transcripts per kilobase million (TPM+1) or fragments per kilobase million (FPKM+1). Then we use the Gene Ontology database [32–34] to select genes that are associated with the following ontologies (hereby referred to as GO genes): “Cell Development”, “Cell Morphology”, “Cell Differentiation”, “Cell Fate” and “Cell Maturation”. Note that the selection of genes still includes a portion of cell cycling genes.

The selected ontologies are ones that are closely related to cell development. We then filter out the selected genes using a dispersion ratio. The dispersion ratio is simply a ratio of the mean over its standard deviation. We set the cut-off to 10 in order to retain at least a few hundred genes. For each gene *j*, we calculate the ratio denoted by ***disp*_*j*_** using the following formula:

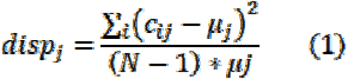

where ***c*_*ij*_** is the j-th gene expression value for the i-th cell and ***μj*** is the average gene expression for the j-th gene over all cells. In all the analyses, the cut-off is set by default to ***disp*_*j*_ ≥ 10** This feature selection procedure reduces the dimensionality problem from ten thousands of genes to a few hundred.

### Consensus Hamiltonian Path (cHMT)

Let X be the gene expression matrix of N cells (rows) by M selected GO genes (columns). We want to infer an N by 1 vector denoting the pseudo time of each cell. This problem can be remodeled as a Hamiltonian path problem with the assumptions of cell similarity within a particular cell type and that the differentiation process is linear. Although many heuristic solutions have been developed to solve this NP-complete problem, most produce inconsistent results due to locally optimal solutions [35]. In order to overcome this difficulty, we developed a consensus Hamiltonian path solution to infer the pseudo time. The algorithm consists of the following main steps:

INPUT: X(N, M)
FOR *k* = 3 to N:

1. X is clustered into *k* groups of cells
2. Generate X’(*k*, M) by taking the average over *k* clusters
3. A Hamiltonian path solution is calculated for each *k*.
Merge each path solution to produce the final solution

Intuitively, we first infer a rough pseudo time ordering of large clusters of cells, then gradually refine the solution with the increase of k.

The clustering method in step 1 is inspired by spectral clustering and SCORPIUS [19]. Briefly, the Spearman correlation distance matrix is first calculated as the pairwise distance between two cell vectors ***x*_*i*_, *x*_*j*_**:

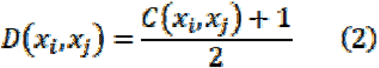

where ***x*_*i*_** and ***x*_*j*_** are both *m*-dimensional vectors of the GO gene expressions and *C()* denotes the Spearman correlation value. Then using the N by N correlation distance matrix, we apply double centering to normalize the matrix. Finally, a simple hierarchical clustering is applied for ***k* ∈ [3… *N*]**.

Next, a Hamiltonian path problem is solved for each value of *k*. In this context, the solution is defined as a path that visits each cell cluster or cell while minimizing the total distance. Hence the definition of the distance function is important to the final solution. A naïve cost function would be to use the Euclidean distance between the cluster centers. However, in order to better model the biological mechanism, we proposed an asymmetric distance measure, namely the Kullback-Leibler distance (KL-distance), or more often referred to as KL-divergence [36]. KL-divergence simply measures the difference between two distributions. In this scenario, we have a pairwise comparison between each *m*-dimensional cell vector: ***x*_*i*_** (i.e., when *k* = N) or cluster averaged vector (i.e., when *k* < N). In the following notations, cell vector will encompass the cluster averaged vector in general.

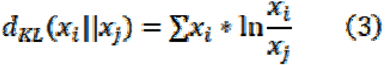

Equation 3 shows the calculation of the KL-distance for two different distributions, namely, from ***x*_*i*_** to ***x*_*j*_**, with each representing an *m*-dimensional distribution of gene expression. The vice versa direction would result in a different value. The intuition here is that the differentiation process is directional and irreversible. Hence, given a more differentiated cell, the distance for it to reverse back to a less differentiated cell should be penalized. Although we cannot be sure of which cell is more or less differentiated, the KL-distance metric gives a small directional restriction in this sense. Direct comparison of the performance of KL-distance and Euclidean distance is given in Figure S1.

After calculating the pairwise distance between each cell vector, we now turn to the modeling of a Hamiltonian path problem. We first define the problem as a graph G = (V, E) where V is the list of vertices, and E represents the number of edges. In our case, each vertex corresponds to a cell vector ***x*_*i*_ = (*x*_*i1*_, *x*_*i2*_,…, *x*_*iM*_)** and the edge weight between each vertex is calculated by ***d*_*KL*_(*x*_*i*_‖*x*_*j*_)** and ***d*_*KL*_(*x*_*j*_‖*x*_*i*_)** as defined in Equation 3. The goal here is to find the shortest path that visits each vertex or cell once.

Next, we developed an O(n^2^)-time heuristic algorithm with a simple modification for the arbitrary insertion algorithm [35] for the Hamiltonian path problem. It should be noted that the Hamiltonian path problem is a classic NP-hard problem, so no algorithm guarantees an optimal solution in any case. Briefly, our heuristic algorithm considers the asymmetric property of our data and modifies the calculation of the insertion cost in the arbitrary insertion algorithm. The main structure of the algorithm remains the same, but we provide a new perspective in solving a Hamiltonian path problem using asymmetrical distances directly. Details of the modified algorithm can be found in the supplementary material. The modified algorithm is performed multiple times, ensuring the quality and robustness of each Hamiltonian path solution. Additionally, a novel approach to find the initial start and endpoints is developed which increased the probability of finding the optimal global solution by the heuristic.

In order to overcome the instability of the Hamiltonian path solutions as well as refining cell heterogeneity at single-cell resolution, we propose a consensus Hamiltonian path in a similar way to reCAT. An advantage of the proposed algorithm is that it will automatically discover the start and end cells of the path. First, a reference Hamiltonian path is built using the enumerated results of five different paths obtained from *k* = 3 to 7 (by default). Since there are two possible starting points of each solution, we then calculate the pairwise correlation between each of the four paths and their respective reverse, yielding ^5^C_2_ = 10 additive correlation scores and 2^5^ = 32 total comparisons. The ordering direction of the best combinations determined by the best correlation score is taken. All five paths are merged to give a reference path by projecting onto the space of [0, 1] and taking the average. The base path is then normalized by feature scaling once again. Hence the direction of the path is determined by the reference path. Subsequently, for each of the following Hamiltonian paths, *k* = 7 to N, the Spearman correlation of each path and its reverse is compared with the reference path. Finally, after adjusting the direction, each path is then merged to the reference path to obtain our final pseudo time. The consensus Hamiltonian path ordering is essentially obtained by sorting the pseudo time values.

Furthermore, we developed an approach to visualize trajectory (linear or branched) development of cells. Intuitively, redPATH recovers a linear pseudo time by merging multiple path solutions to a single path, which naturally merges branching situations as well. We hypothesize that the branching trajectory can be detected by observing the detail transition of cells in each of the merged solutions.

The main idea is to transform multiple Hamiltonian path solutions into a transition probability matrix followed by PCA visualization. Given *p* Hamiltonian path solutions, we record all the transitions in each path *pi* and construct an N by N transition matrix. For instance, if N = 3 and the *pi*-th solution is 2-3-1, then we add a probability value for the transitions 2–3 and 3–1. The transition probabilities are added together until all Hamiltonian path solutions are recorded.

### Biological Analysis

With the Hamiltonian path ordering, we can identify key genes or gene modules specific to the differentiation process. In order to quantify the expression changes over the pseudo time series, we used two statistical measures: maximal information coefficient (MIC) and distance correlation (dCor) [37,38]. Compared to the standard Pearson or Spearman correlation coefficients, these measures are more robust and have a range of [0, 1]. The scores are calculated for all of the genes and ranked accordingly. The genes which exceed a threshold of 0.5 are selected for downstream analysis. However, the downside of these measures is that they cannot determine a positive or negative correlation.

Then we designed a simple hidden Markov model (HMM) to infer two (or possibly three) hidden states of each gene. The two hidden states represent an on/highly expressed or off/lowly expressed state in each cell. The observed variable is simply the gene expression value. In this model, we assume a univariate Gaussian distribution over the two hidden states.

The model is initialized with ***N*[*μ*_*n1*_,*a*_*n1*_]** and ***N*[*μ*_*n2*_,*a*_*n2*_]** where the mean and standard deviation are estimated with the sorted observed gene expression values. Then the transition probability is inferred using the Baum-Welch algorithm [39]. Subsequently, the Viterbi algorithm is implemented to infer the hidden states of the gene in each cell. The inferred states are then clustered using hierarchical clustering and visualized through a heatmap to provide an overall understanding of the gene expression changes over the developmental process. The gene clusters are further analyzed using GOsummaries [40] and PANTHER [41], which provides some biological insights to the gene modules.

### Coupling the differentiation and cell cycle process

In order to identify the relationship between cell differentiation and proliferation, we incorporate reCAT and redPATH to visualize their relationship. One of the challenges faced in analyzing the cell cycle is the removal of G0 cells. To date, there is currently no known algorithm to identify G0 cells. Here, we developed a novel approach using statistical tests to identify the G0 cells before continuing further analysis.

The intuition for the developed approach is that G0 cells tend to be inactive in terms of cell cycling genes, and they are in a resting phase. Hence, we hypothesize that G0-like cells will have the lowest cycling expression. We first transform the gene expression matrix into average expression values for each of the following six mean cell cycle scores, G1, S, G1/S, G2, M, G2/M. This is adapted from reCAT, and we used the annotation from Cyclebase to calculate the average scores. Then we apply k-means with k set to 5 (i.e., G0, G1, S, G2, M stages) to the mean scores. Pairwise analysis of variance (ANOVA) tests were performed for each of the mean scores for the group that was least expressed. The criterion is set such that the identified group must be significant (p-value < 0.001) in all of the six mean scores in its comparison with the remaining groups. The results are validated on a couple of datasets where the G0 cells are known (Figure S2).

After the removal of G0 cells, we inferred the pseudo differentiation and cycling time for each cell using reCAT and redPATH, respectively. Then we produced 3-D spiral plots as an attempt to visualize their relationship. Briefly, the pseudo time of reCAT is projected onto a circle as the X and Y axis, and then the differentiation time is plotted on the Z-axis. Marker genes are used to depict the gradual change of the cell types in each dataset.

### Evaluation Metrics

In order to quantitatively assess the pseudo temporal ordering, we used four metrics to compare our results with existing algorithms. There are limitations to evaluating the accuracy of the orderings because the delicate ordering within each different cell type remains unknown. The only information available is the cell type labeling obtained from biological experiments, which may also potentially contain some bias due to technical noise during biological experiments. Using the cell type information, we developed change index (CI), bubble sort index (BSI), and further applied Kendall correlation (KC) and pseudo-temporal ordering (POS) score to evaluate the reconstructed pseudo-time.

For illustration purposes, an example of a linear development of different cell types is used here. A linear development of the neural system in the sub-ventricular zone is defined from quiescent neural stem cells (qNSC) to activated neural stem cells (aNSC) then differentiating into neural progenitor cells (NPC). In other words, we assume that we have 3 stages of development, ordering from qNSC to aNSC to NPC.

The first metric, change index, was adopted from reCAT [20]. Assuming the number of states is *ns* (which is 3 in our example), we calculate the number of state changes, *s*, after re-ordering the cells. Then we calculate the change index as CI = 1 − (*s* − *ns* − 1) / (N − *ns*) where N is the total number of cells. Hence, a temporal ordering that completely resembles the true labeling of cell types would have a value of 1 and the worst case of 0.

From experimental results, we found that the change index may be inaccurate when a large subset of a particular cell type is grouped together, but inserted within another cell type of development. Hence we designed a second metric called the bubble sort index to evaluate the re-ordered time series. The intuition behind this index is inspired by the number of steps, *s*, taken to re-sort the time series. This is basically the number of moves of switching adjacent cells that it needs to make to correct the ordering and has better stability over the change index. The number of steps *s* is then divided by S, which is the number of steps taken to sort the worst-case scenario (i.e., the reverse of the correct ordering), to produce the bubble sort index. Generally, the bubble sort index results in higher values in the range of [0, 1].

Thirdly, we also used the Kendall correlation coefficient to evaluate our time series. Both Spearman and Kendall correlation would work better than the Pearson correlation in this case due to the consideration of ranking in the implementation of these two methods. Additionally, the POS score is also adapted from TSCAN [18] to evaluate the performance of each algorithm.

## Results

### Validation and evaluation of redPATH

#### Introduction

The intuition of redPATH is first validated, and its performance is then compared with current state-of-the-art algorithms. This comparison is mainly based on three neural stem cell datasets [6,7,42], one hematopoietic dataset [8], one human hematopoietic dataset [43], and three embryonic time point datasets [44–46]. The further downstream analysis included recent glioma datasets [5] to uncover underlying mechanisms behind cancerous cells. All the datasets used are listed in **Table 1**.

**Table 1.** Description of analyzed datasets

| **Table 1. Description of analyzed datasets** |  |  |  |
| --- | --- | --- | --- |
| **Datasets** | **No. of cells** | **No. of cell types** | **Organism** |
| Neural Stem Cells: |  |  |  |
| NSC-Dulken (PRJNA324289) | 250 | 3 | *Mus musculus* |
| NSC-Llorens-B (GSE67833) | 145 | 3 | *Mus musculus* |
| NSC-Shin (GSE71485) | 168 | 2 | *Mus musculus* |
| Hematopoietic Cells: |  |  |  |
| HC-Schlitzer (GSE60783) | 251 | 3 | *Mus musculus* |
| Embryonic Stem Cells: |  |  |  |
| mESC-Deng (GSE45719) | 268 | 10 | *Mus musculus* |
| mCV (GSE100471) | 598 | 3 | *Mus musculus* |
| mGas (GSE100597) | 639 | 4 | *Mus musculus* |
| Glioma Cells (GSE89567): |  |  |  |
| MGH45 – WHO IV | 594 | 4 | *Homo sapiens* |
| MGH57 – WHO IV | 334 | 1 | *Homo sapiens* |
| MGH107 – WHO II | 252 | 3 | *Homo sapiens* |
| Human HSC (hHSC): (GSE70245) | 382 | 4 | *Homo sapiens* |

For the neuronal dataset of Dulken and Llorens-Bobadilla, both studies look at the development of neural stem cells (NSCs) in the subventricular zone (SVZ), whereas Shin’s data was obtained from the subgranular zone (SGZ). The development lineage is quite clear where quiescent NSC (qNSC) becomes activated NSC (aNSC) and further differentiates into neural progenitor cells (NPC) and finally into neuroblasts (NB) or neurons. The hematopoietic data looks at the development of dendritic cells near the end of the lineage. The macrophage and dendritic cell precursor (MDP) differentiate into common dendritic cell precursors (CDP) and give rise to pre-dendritic cells (preDC). An important question of interest is how differentiation and proliferation processes are regulated within these different cells. This is explored in the later parts of this paper, which discusses the incorporation of reCAT and redPATH to provide a simple exploratory analysis.

#### Quantitative evaluation of redPATH

First of all, the modeling of single-cell trajectory as a Hamiltonian path problem needs to be confirmed as a valid approach. From **Figure 2A**, we can see that the developmental process across cells is aligned by the Hamiltonian path for *k*=3 and *k*=7 clusters. This sets the foundation for redPATH. Assuming the order of the development progression is correct, the ordering is refined by combining the paths of larger *k* and thereby obtaining a stable solution.

**Figure 2.**
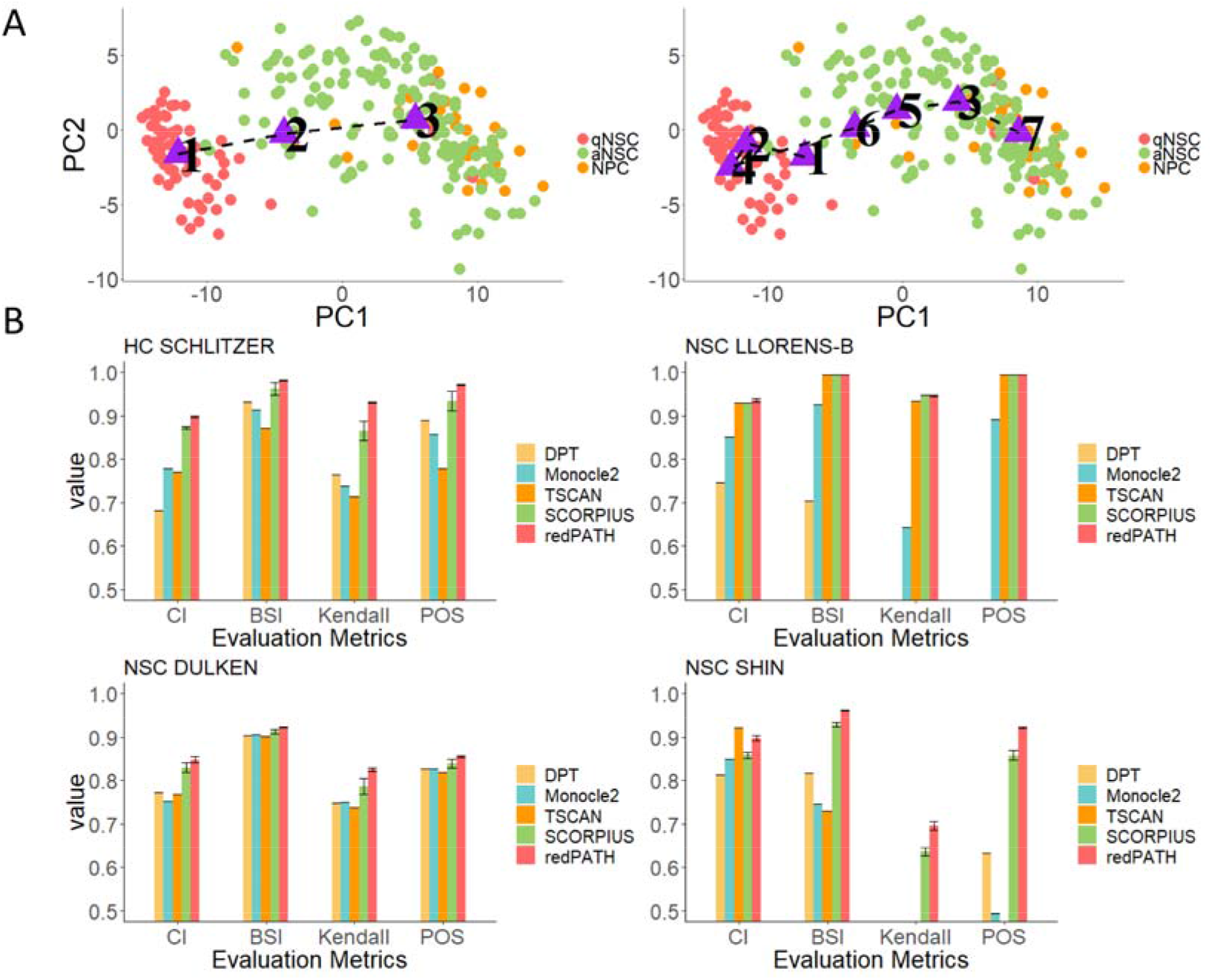
Validation of redPATH. **A** shows the Hamiltonian path solution by projecting each cell and cluster centres onto scaled and centred principal components (PCs) 1-2 for k=3 and k=7 respectively. The purple triangle represents the cluster centres and dotted line reflects the Hamiltonian path. Each cell is colored by its cell type label. **B** provides the performance evaluation of different algorithms on four single-cell datasets using change index (CI), bubblesort index (BSI), Kendall correlation (KC), and pseudo-temporal ordering score (POS). Bar plots are colored by the algorithm used, and the rightmost bar (in red) represents redPATH. The error bar shown represents the 99% confidence interval based on 20 runs of the algorithm. Missing bars in the plots represent an evaluation value of less than 0.5.

redPATH is compared with Monocle2, TSCAN, and SCORPIUS for its performance. The results are shown in **Figure 2B**, where redPATH consistently shows the best performance across all the scores for the three neuronal datasets and one hematopoietic dataset.

A comparison is made by using the same input (the selected Gene Ontology genes) for each of the algorithms. SCORPIUS claims to be robust when using all the genes without gene selection, but the performance did drop by a small margin across all datasets when using the full gene expression matrix. It should be noted that NSC-Llorens-B performed quite well overall partially because the data was sequenced at a much deeper length. The rightmost bar from **Figure 2B** represents the redPATH method. The error bar represents a 99% confidence interval based on 20 runs of both SCORPIUS and our algorithm. Furthermore, redPATH (CI: 0.69, BSI: 0.92, KC: 0.82) is on par with SCORPIUS (CI: 0.62, BSI: 0.92, KC: 0.84) on a multi-time point dataset (mESC – Deng) with ten cell types. The outperformance of the change index also proves its capability to analyze time point data as well as snapshot data. Additional multiple time-point datasets are evaluated, and results are shown in the supplementary Figure S3.

The performance of many algorithms may be susceptible to cell subpopulation and different gene selections. In **Figure 3**, we present the robustness of each algorithm on subsamples of cells. For each of Llorens-B-NSC and Dulken-NSC datasets, we sampled 30%, 50%, 70%, and 100% of all cells 20 times. As shown in **Figure 3A**, the evaluation of redPATH on all three metrics is relatively consistent and stable; a similar pattern is observed in **Figure 3B**. A comparison of the gene feature selection approach is also included in the supplementary Figure S4. Additionally, we also compared performance of redPATH with a different set of selected genes using dpFeature (Figure S5).

**Figure 3.**
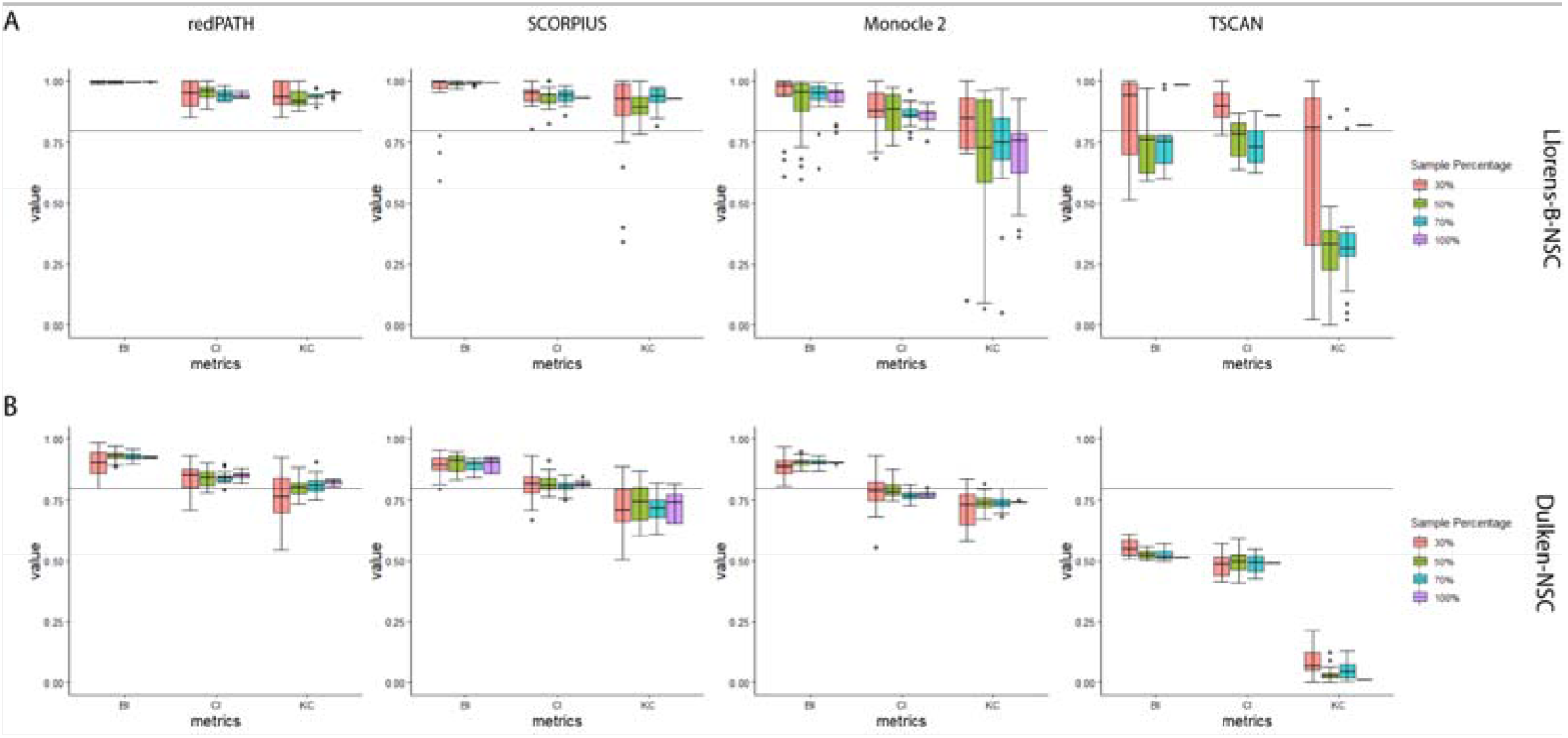
Robustness analysis on algorithms. **A-B** 30%, 50%, 70%, and all cells are sampled from the Llorens-NSC and Dulken-NSC datasets, respectively, and each column represents different algorithms. Each algorithm is run for the same 20 subsamples and is evaluated on BI, CI, and KC. The boxplot represents the standard quantile range for the calculated values. The horizontal line denotes the 0.8 mark for the evaluation value.

#### Observing the differences in inferred biological development

Accounting for all the metrics across each dataset, SCORPIUS has a relatively better performance than the rest of the other methods. In order to further explore the differences in biological functions between redPATH and SCORPIUS pseudo-time, we observe the developmental trend on some marker genes on all three NSC datasets (**Figure 4**). *Stmn1* and *Aldoc* [42,47,48] are considered to be marker genes for the differentiation of neural stem cells. *In vivo* experiments [42] had been conducted to show that *Stmn1* is highly expressed in NPC with little activity in NSC, and *Aldoc* is only expressed in quiescent NSCs and low-expressed in aNSC and NPC. We compared gene expression development for redPATH and SCORPIUS due to the overall better performance of these two algorithms (**Figure 4**). A comparison of additional marker genes is included in the supplementary (Figure S6).

**Figure 4.**
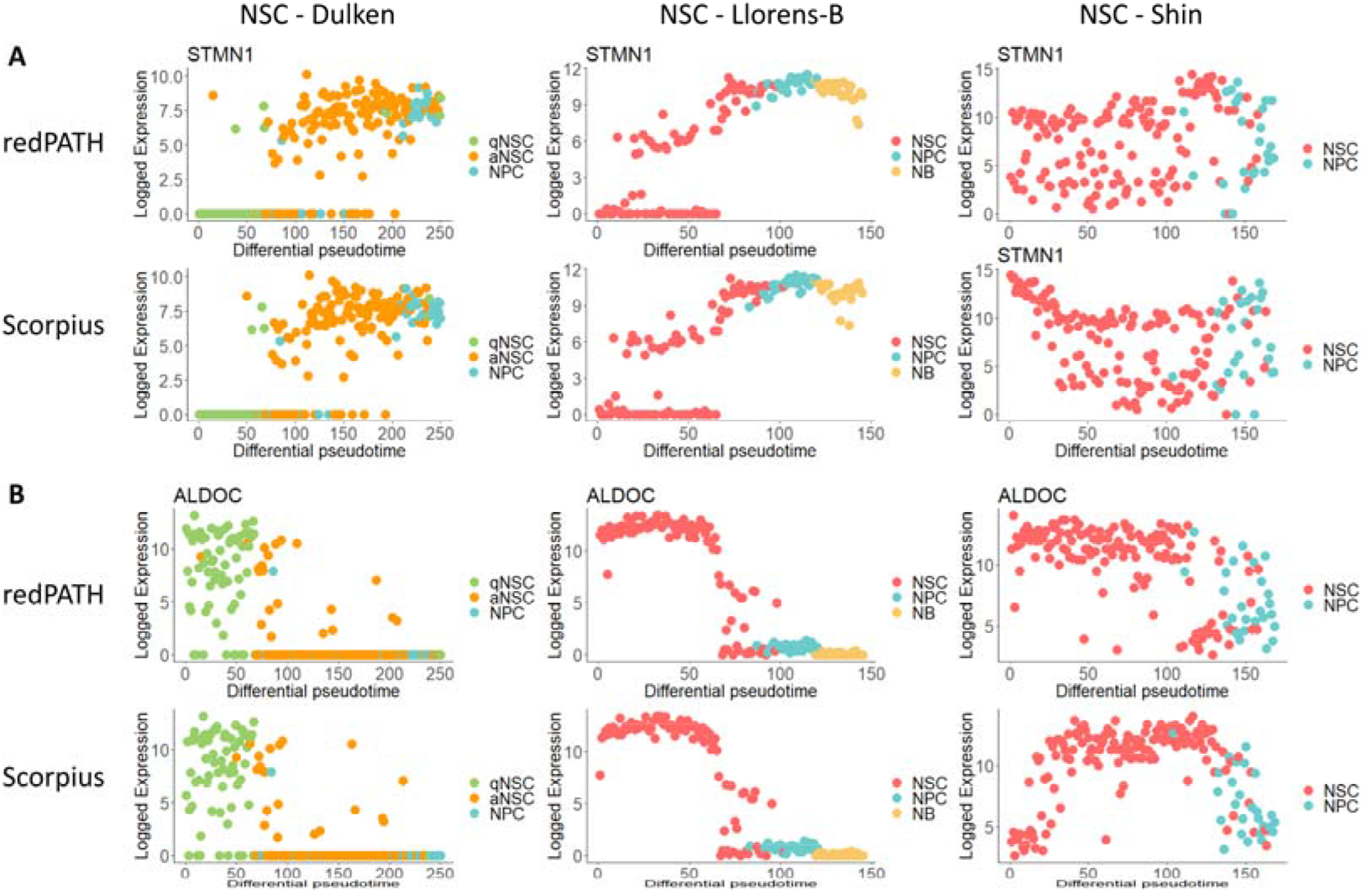
Qualitative comparison on expression changes. **A** depicts the difference in gene expression trend for *Stmn1* by plotting the gene expression against inferred pseudo time. Comparison is made across three NSC datasets (Dulken, Llorens-B, and Shin respectively for each column) using redPATH and SCORPIUS. **B** Similarly for *Aldoc*.

On both NSC-Dulken and NSC-Llorens-B dataset, the performance of redPATH is on par with SCORPIUS, and no significant difference is observed. In the rightmost panel (NSC-Shin dataset), the ordering of SCORPIUS clearly shows a different patterning compared to the other NSC datasets. With the *Stmn1* gene (**Figure 4A**), SCORPIUS starts with a high expression (which is supposed to be lowly expressed at the start of the trajectory), then decreases, which is different from the conclusion made from biological experiments. redPATH fits the developmental trend with relatively low expression at the beginning of the trajectory and shows consistency across datasets for the same cell type. We can observe that SCORPIUS tends to identify some bell-shaped trend, which could be explained by iteratively fitting principal curves in their algorithm. This observation can also be made from **Figure 4B** in the NSC-Shin dataset. Here, redPATH proves to be robust across different datasets and correctly orders the developmental pseudo-time in accordance with biological observations.

#### Identifying trajectory development of cells

Utilizing the multiple Hamiltonian path solutions from redPATH, we can construct a cell transition matrix and visualize the developmental trend on a PCA plot (**Figure 5**). The trajectory plots are shown for two linear progression datasets, Llorens-B-NSC and Dulken-NSC, as well as a branching hematopoietic stem cell dataset (hHSC). The progression in NSC cells along the pseudo time reflects that there is a linear development from NSC to NPC (**Figure 5A-B**). However, for the hHSC dataset, the PCA plot suggests a branching development of cells (**Figure 5C**), confirming with the original discovery of binary cell fate decisions [49]. There appear to be two separate progressions of cell differentiation. A comparison of trajectory plots produced by different algorithms can be found in the supplementary Figure S7 and S8.

**Figure 5.**
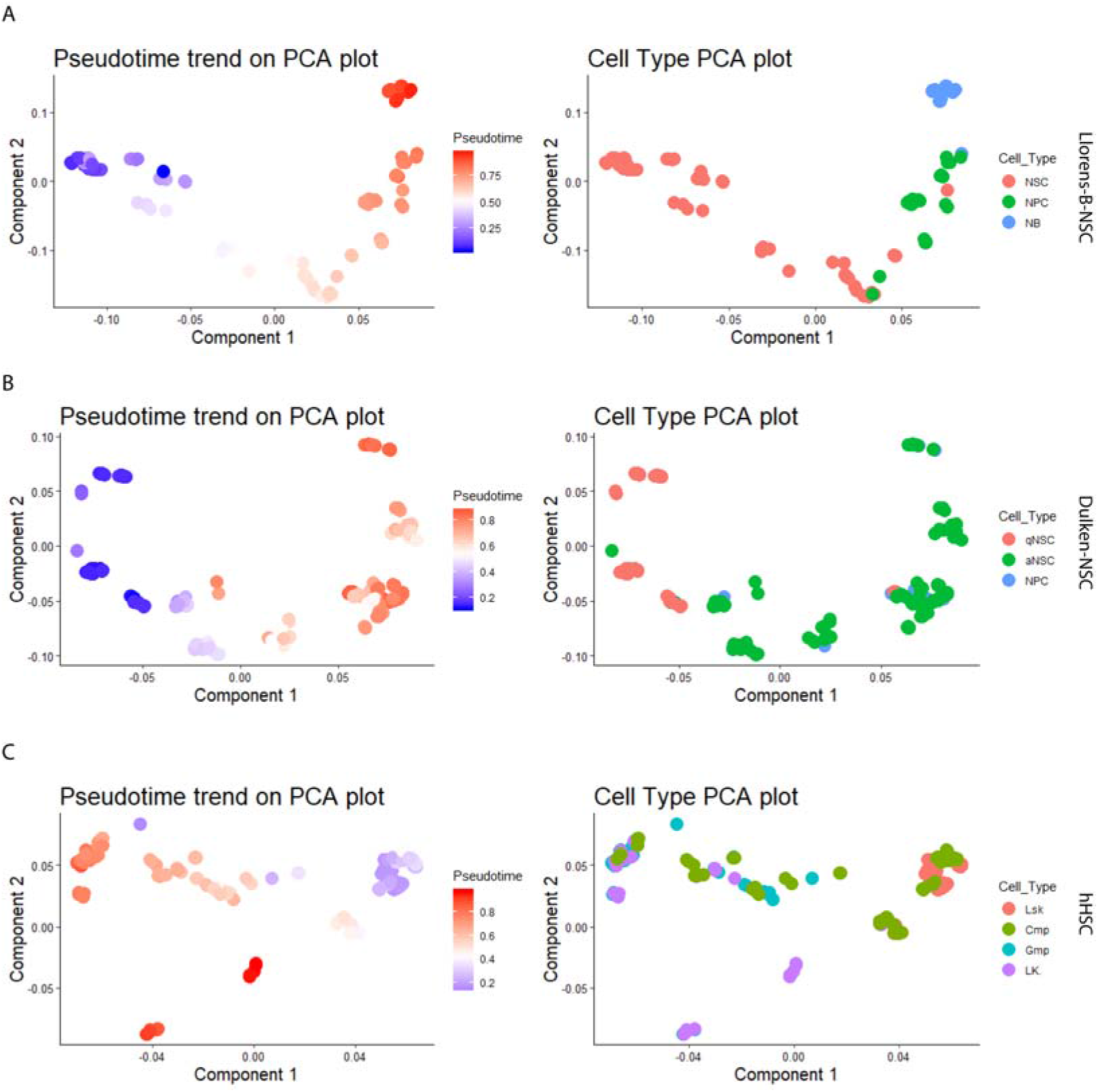
Trajectory development of cells. **A-C** Visualization of the differentiation development process colored by pseudo time and cell type information for Llorens-NSC, Dulken-NSC, and hHSC, respectively. Each point represents a cell in space, and the PCA is performed on the calculated transition matrix. The left panel depicts the pseudo time of each cell, and the right shows the corresponding cell type information.

#### Coupling proliferation with differentiation

As an attempt to visualize the relationship between the cell cycle process and differentiation, 3-D plots are produced for the NSC-Llorens-B dataset. Before analyzing the relationship between cell proliferation and differentiation, G0-like cells are removed from the dataset. The developed approach was run twice to remove all possible G0 cells from the dataset (with a threshold of p-value < 0.001). The differential pseudo-time is re-calculated with redPATH on the remaining cells, and cell cycle analysis results are obtained from running reCAT. Here redPATH (CI: 0.862, BSI: 0.977, KC: 0.852) outperforms SCORPIUS (CI: 0.828, BSI: 0.853, KC: 0.589) on the remaining 61 cells, showing its reliability even in a very small sample dataset. NSC marker genes (*Egfr, Stmn1 [7,42]*) further validates that most G0-like cells have been removed from the downstream analysis, where neither expresses much during the quiescent state (**Figure 6**).

**Figure 6.**
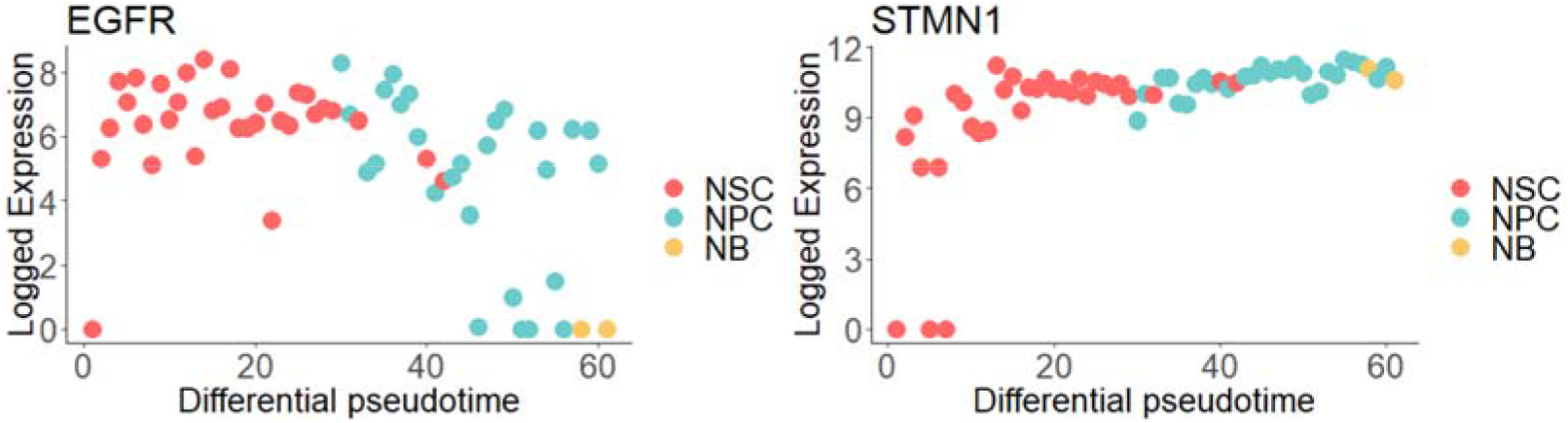
Validation on the removal of G0-like cells. Llorens-NSC is processed by removing G0-like cells, and pseudo time is calculated on the remaining cells. Expression changes of *Egfr* and *Stmn1* are plotted against the inferred differential pseudo time.

Using the two evaluation statistics of distance correlation (dCor) and maximal information coefficient (MIC) at the threshold of 0.65, we uncovered three genes (*Foxm1, Tubb5, Nek2*), which correlates highly with both cell proliferation and differentiation. Differentially associated marker genes such as *Dcx, Dlx1-2, Dlx5, Tubb3, Cd24a, Sox11, Dlx6as1, Mfge8, Sp9,* and *Atp1a2*, are in concordance with previous studies [6,7,47,50]. Similarly, we also uncovered interesting genes that are cell cycle-related. For example, *Cdk1* and *Aurkb* which associate with cell proliferation and NSC activations.

*Foxm1* was recently reported to regulate a micro-RNA network which controls the self-renewal capacity in neural stem cells [51]. redPATH provides an interactive plot that can visualize different cell types, cell cycle stages, and gene expression together. Reducing the left panel of **Figure 7** to NSC and NPCs, *Foxm1* is highly expressed in G1 and G2/M cycling stages, which is indicative of cell proliferation. Observing NSCs (the inner orange points on the left), a subset of cells within the ellipse is lowly expressed as compared to the outer orange points. This could suggest that NSCs may be at its earlier stages of activation, which is more quiescent-like as compared to the higher expressed activated NSCs.

**Figure 7.**
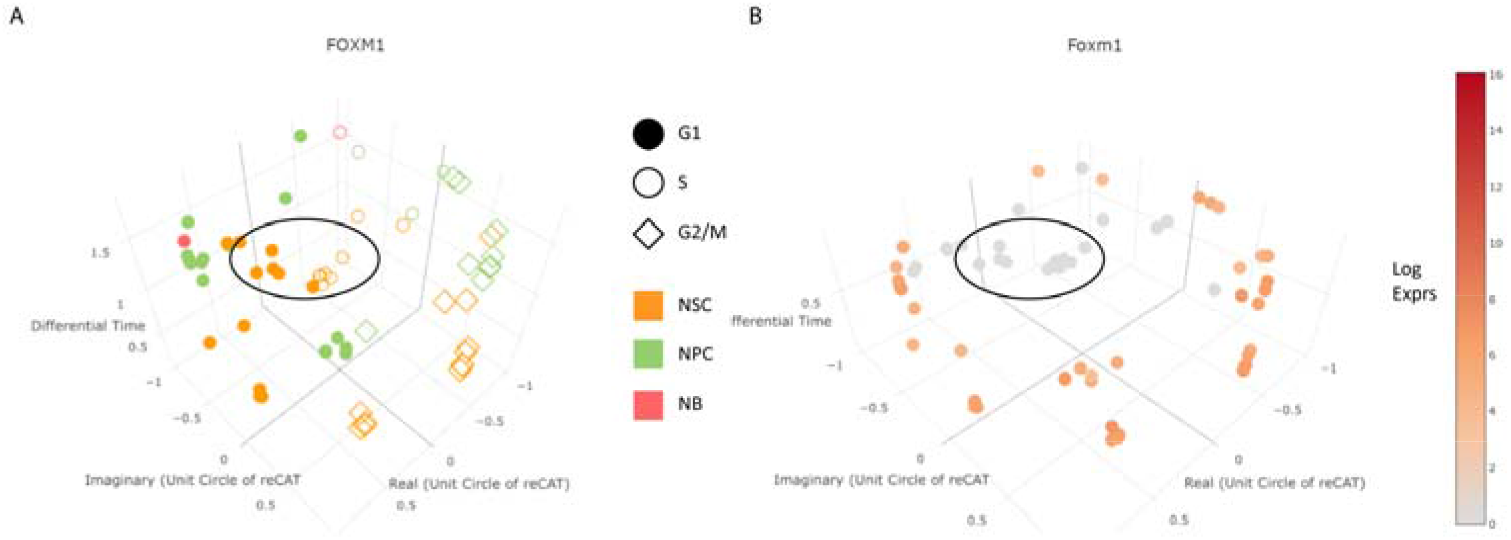
3D visualization of cell proliferation and differentiation on *Foxm1*. **A** plots the differential pseudo time (z-axis) against proliferation pseudo time (x- and y-axis) colored by cell type and cell cycle stages. **B** shows the gradual change in expression for the 3D plot.

#### redPATH analysis on glioma datasets

Assuming that the snapshot on the cancerous dataset provides the different development stages of single cells among the dissected tissue, we can uncover some underlying mechanisms by observing the pseudo temporal development of gene expression change from microglia/microphage cells to malignant cells within a tumor dissection. Normal microglia cells exist to eliminate any intruding cells, also acting as antigen-presenting cells which activate T-cells [52]. However, immune functions of microglia/macrophage cells within glioma tumors are impaired and are more commonly known as glioma-associated microglia/macrophages (GAMs), which regulate tumor growth [9,10,12,13]. As the original publication [5] suggests, malignant cells include some properties of neural stem cells with active differentiation in glial cells specifically. Although the tumor microenvironment is much more complicated, gene modules and possible relevant genes can be inferred.

#### Gene module extraction

In the original publication [5], the authors have classified each tumor cell as either malignant cell, microglia/macrophage, oligodendrocyte, or T cell using clustering and copy number variation analysis. Although these four cell types do not differentiate into one another, GAMs and T cells are altered to regulate malignant cells. Here, we re-ordered the cells using redPATH and successfully recovered a pseudo developmental trend to observe gene expression change.

MGH107, a grade II astrocytoma that has not been treated yet, shows a gradual change in gene expressions indicating a subpopulation of malignant cells. The other two grade IV tumors showed less progression but still revealed a subpopulation in MGH57 (Supplementary Figure S9).

Using dCor and MIC, 921; 55; 762 significantly identified genes are retained for analysis for MGH45, MGH57, MGH107 respectively (threshold >= 0.5). The gene expression profile of oligodendrocytes is closer to malignant cells. Here, the result for MGH107 is shown (**Figure 8**).

**Figure 8.**
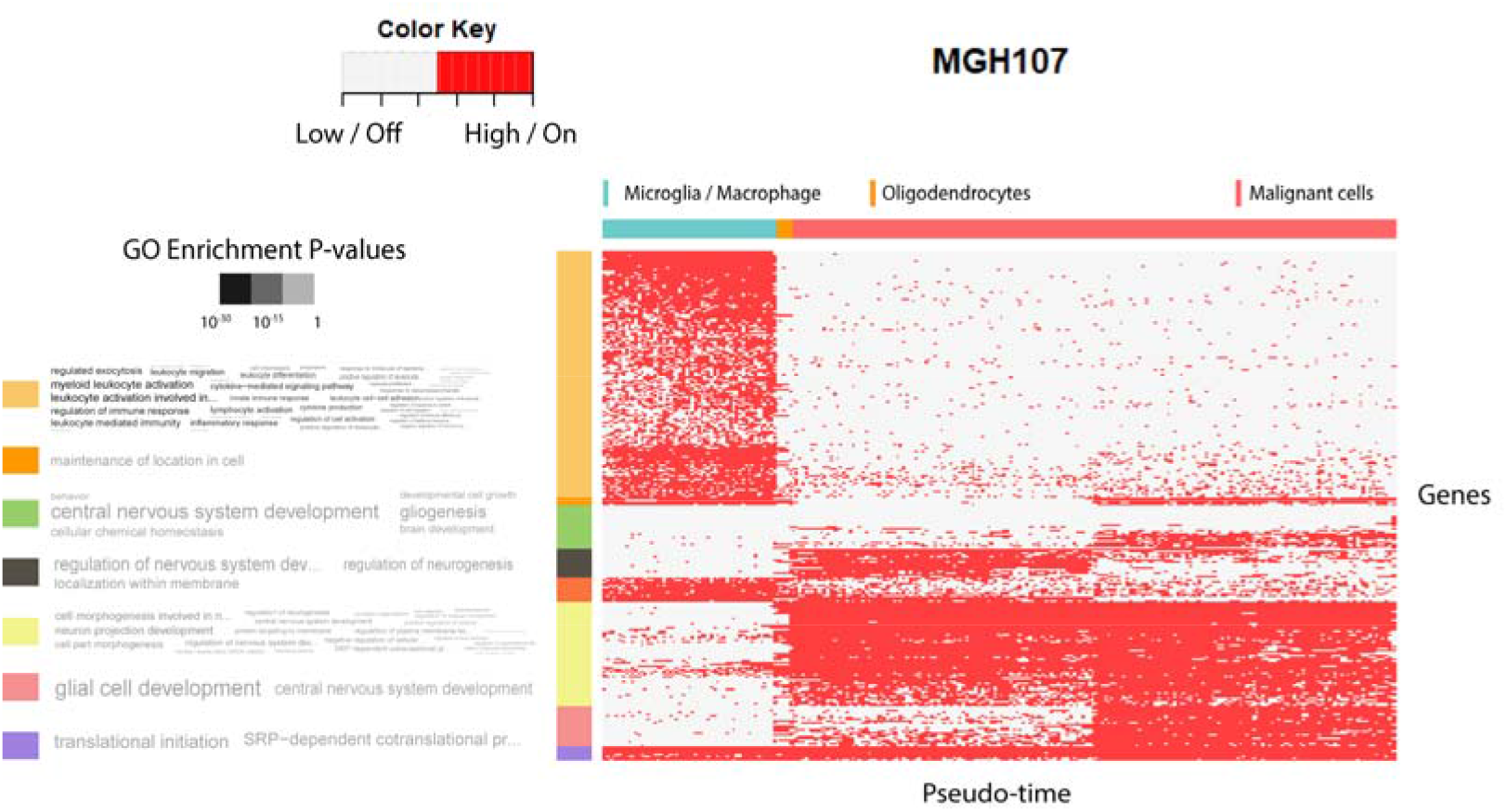
Heatmap analysis on MGH107. Heatmap plot is produced according to the inferred HMM results from redPATH, indicating on / highly expressed state or off / lowly expressed state of each gene. The horizontal ordering denotes the differential pseudo time while each row represents a significantly identified gene. Gene clustering is shown on the left with Gene Ontology enrichments.

#### Stem-cell like subpopulation in glioma cells

Focussing on the glial cell development / the central nervous system development gene module of MGH107 in **Figure 8**, astrocytic and stem cell-like markers (*ATP1A2, GFAP, CLU, ALDOC [5,42,53]*) are found to be expressed in the latter half of the malignant cells while quiescent markers such as *ID3* remained silenced. Additionally, a subpopulation of malignant cells can be clearly identified by observing the top-ranked identified genes such as *VIM, SPARCL1, TIMP3* (Supplementary Figure S5). This indicates a high potency of the malignant cells to differentiate and proliferate. The malignant cells of Grade IV glioblastoma (recurrent) MGH45 show a constant gene expression pattern. However, MGH57 (Grade IV glioblastoma) revealed a relatively small subpopulation of malignant cells that does not express *OLIG1, OLIG2, DLL1, CCND1, IGFBPL1,* and express *ALDOC* and *ATP1A2* (Supplementary Figure S9). Here, *ATP1A2*, *IGFBPL1,* and *ALDOC* are all possible significant stem-like markers from prior analysis on the neural stem cells mentioned above. These results indicate a subset of non-proliferative malignant cells in MGH57 and MGH107. MGH45 is a recurrent glioblastoma patient, hence it is possible that a large portion of malignant cells are stem-cell-like.

#### Apoptosis program within different gliomas

An interesting exploratory finding is the apoptosis program within gliomas. Apoptosis is a mechanism within the body that is activated intrinsically or extrinsically which leads to cell death. All three tumor patients had not been treated with medication or radiation before; hence external factors of cell death are not applicable.

*MCL1 [54–56]*, an important BCL-2 family apoptosis regulator is significantly expressed within the same gene cluster of “glial cell development” (dCor: 0.59, MIC: 0.50). The expression of *MCL1* activates *BAX* and *BAK* modules in the apoptosis pathway in general. Also, it has been recently reported [56] that silencing *MCL1* leads to inhibition of cell proliferation, thereby promoting apoptosis in glioma cells. Here, it can be observed in **Figure 9A** that there are two subpopulations for the malignant cells expressing in *MCL1*. **Figure 9A** of MGH45 also shows that microglia are inhibited. The proportion of malignant cells, which possibly promotes apoptosis to proliferating malignant cells, are similar: MGH107 - 0.45, MGH57 - 0.5, MGH45 - 0.35.

**Figure 9.**
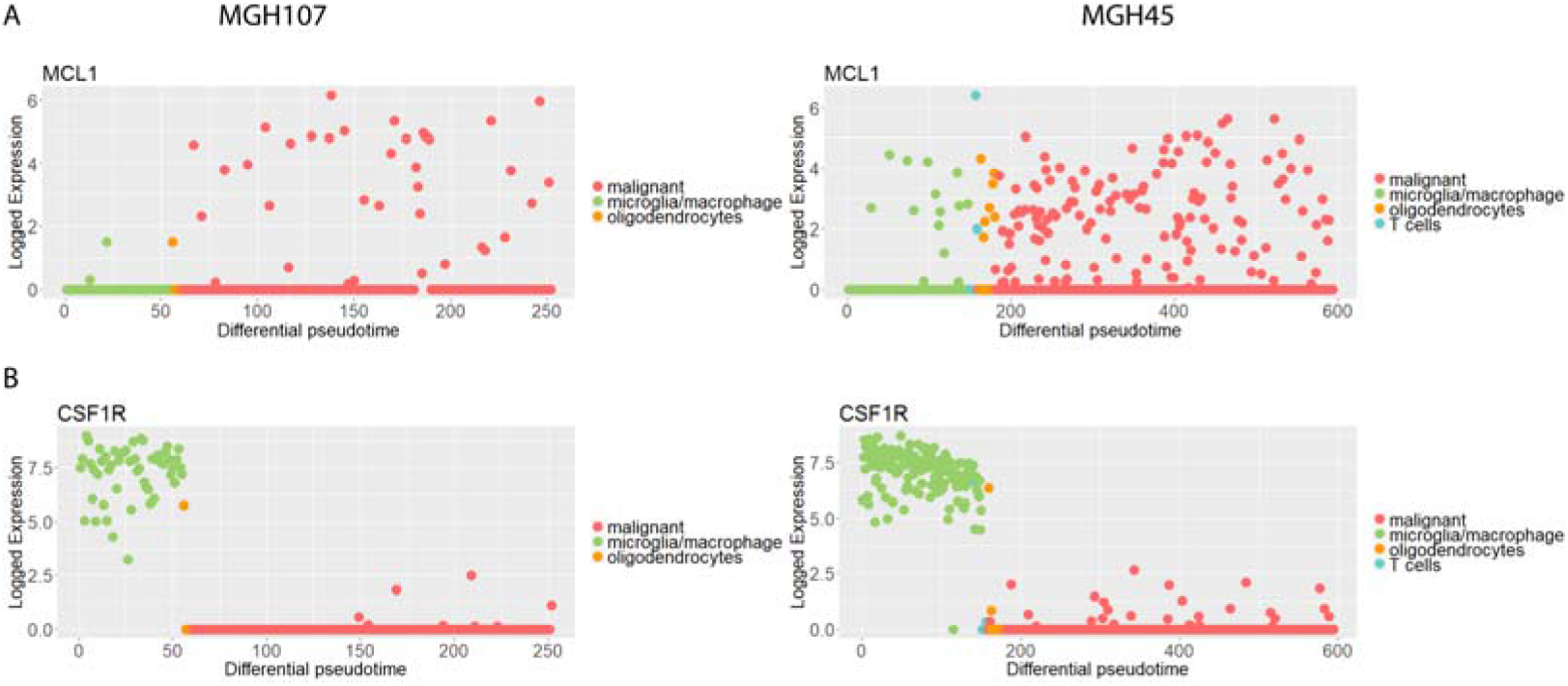
MCL1 and CSF1R expression changes in glioma cells. **A** shows the expression trend along the pseudo time in *MCL1* for MGH107 and MGH45, respectively. Similarly for *CSF1R* in **B**.

Intuitively, the situation of MGH45 appears to be quite severe, where only a small number of cells activate apoptosis. Although numerous other apoptosis signaling pathways are available, further biological validation would be beyond the scope of this analysis. Drugs targeted at the BCL-2 family and MCL-1 inhibitor was under pre-clinical trials in 2015 with promising results [11,57,58].

#### Discovery of potential significant genes

Additionally, we ranked the top genes in the supplementary. There are numerous overlaps in MGH45 and MGH107, where *CSF1R* (dCor: 0.95; 0.93, MIC: 0.78; 0.78) is discovered with distinct change between microglia / macrophage and malignant cells. It has been previously reported that inhibition of *CSF1R* in macrophages may lead to a re-programming of macrophages, which in turn reduces tumor growth [14,59]. However, experiments also showed that inhibition of *CSF1R* eventually acquires resistance and *PI3K* signaling pathways are activated to support malignant cells [15]. It is trivial from **Figure 9B** that the microglia/macrophages are overly expressed within the tumor microenvironment. Additional marker genes can be found in the supplementary (Supplementary Figure S10).

Overall, redPATH can be utilized to analyze single-cell transcriptome datasets with and without cell type labeling. As shown in the heatmap analysis of glioma cells, redPATH can also correctly recover the cell type segmentation along a developmental pseudo-time.

## Discussion

With the initial intent to analyze pseudo developmental processes of single-cell transcriptome data, we developed a novel comprehensive tool named redPATH to provide computational analytics for understanding cell development as well as cancer mechanisms. redPATH shows its robustness in recovering the pseudo-developmental time of cells and its capability in detecting both branched or linear progressions. The algorithm demonstrates high consistency across different sample numbers as well as different feature selection methods. Subsequently, analytical functions implemented include: 1) detection of G0-like cells, 2) gene discovery using dCor and MIC, 3) 2- or 3-state HMM segmentation inferring low / highly expressed gene state, 4) gene module extraction and 3D visualizations for differentiation and proliferation processes, and 5) visualization for identifying branched or linear cell development.

In this manuscript, we show that redPATH is capable of recovering the cell developmental processes successfully and we analyze glioma datasets with a new perspective. This results in the discovery of stem-cell-like and apoptotic marker genes (such as *ATP1A2, MCL1, IGFBPL1, ALDOC*) along with a deepened understanding of diseases and cell development. It is capable of discovering significant novel genes using the pseudo-time rather than testing the differential genes by groups. Although the advantage is that cell type labeling is not required here, this approach may fail when the pseudo-time results perform poorly.

redPATH attempts to visualize the coupling relationship between cell proliferation and differentiation; however, integrative models are preferred to analyze such processes simultaneously. The underlying mechanism remains obscure and requires more integrative computational models. Furthermore, biological validations are required for the identified lists of significant genes.

## Availability

redPATH is available through https://github.com/tinglab/redPATH. Details about the data used in this manuscript can be accessed in **Table 1**. Briefly, data accession numbers for neural stem cells include: PRJNA324289, GSE67833, GSE71485; hematopoietic cells: GSE60783, GSE70245; embryonic stem cells: GSE45719, GSE100471, GSE100597; and glioma cells: GSE89567.

## Authors’ contributions

KX, ZL, NC, and TC designed the study. NC and TC supervised the study. KX carried out the main implementations and data analysis. ZL provided interpretations in the cell cycle analysis. KX, ZL, NC and TC wrote the manuscript. All authors read and approved the final manuscript.

## Acknowledgements

This work is supported by the National Natural Science Foundation of China (grants 61872218, 61721003, and 61673241), Beijing National Research Center for Information Science and Technology (BNRist), and Tsinghua University-Peking Union Medical College Hospital Initiative Scientific Research Program. The funders had no roles in study design, data collection and analysis, the decision to publish, and preparation of the manuscript.

Thank Zehua Liu and Ting Chen for fruitful discussions on this work. Thank Professor Ron Shamir for the inspiration of the bubble sort index. We also thank David Pellow, Zhen Zhong and Israel Goytom Birhane on discussions about cell development. Grateful to Professor Jon Rittenhouse, Louisa Zhang, and Tong Lee Chung for their help on proofreading and editing; Yanan Ru and Xiaoqing Su for suggestions on visualization themes.

## Supplementary material

It has been uploaded as a separate file: Supplementary Materials S1.

