## Supplementary File S1 for "redPATH: Reconstructing the Pseudo Development Time of Cell Lineages in Single-Cell RNA-Seq Data and Applications in Cancer"

***Modeling the cell differentiation by asymmetric distance (KL-distance)***

We proposed to model the cell differentiation process by using KL-distance combined with consensus Hamiltonian path algorithm. The intuition is that the asymmetrical property of KL better reflects the direction of differentiation development, we believe that it can capture the biological information better. A comparison between the Euclidean distance and KL distance is made on subset data from four single-cell datasets (Figure S1). We sampled 8 cells, 20% of the cells, and 80% of the cells 50 times and conducted the evaluation. Brute force search is performed on subset of 8 cells, and redPATH is performed on the other two subsets.

On the left panel of Figure S1A, it is quite clear that the shortest path (calculated by brute force search) from KL-distance achieves a significantly better evaluation. In Figure S1 B-D, KL-distance achieves at least as good as Euclidean distance in worst-case. As N increases, the results become more stable across all four datasets.

***Identification of G0-like cells***

To date, there is no existing method to identify G0-like cells. Here we provide a simple statistical test to identify possible G0 cells in different datasets. The results are validated by using NSC-Dulken dataset, where quiescent state NSCs (qNSC) have been labeled. qNSC cells are inactive stem cells which retain the ability to proliferate and differentiate upon activation.

First, we assess the mean scores of each dimension (namely G1, S, G1/S, G2, M, G2/M) of cell cycling genes. Kmeans clustering (K = 5) is performed on the 6 dimensional mean scores to separate clusters of cells. The distribution of each cluster is shown in Figure S1A.

By applying a simple ANOVA test to each pairwise comparison, we identified that group 4 (Figure S2A) as possible G0 cells with a threshold of p-value < 0.001 across all comparisons. Figure S1B confirms our approach as most of the qNSC cells are included in this cluster, with little cells from aNSC and NPC. A small portion of aNSC are identified possibly due to the cell being in its earlier stages of activation. Neural progenitor cells (NPC) contained both proliferative and in-active cells, here, only 3 / 29 were identified as G0.

This procedure is repeated until one or more ANOVA tests become insignificant. The NSC-Llorens data analyzed in the main manuscript was performed twice to eliminate most of the G0-like cells.

***Comparison on feature (gene) selection***

Different gene selections will produce different results, especially for algorithms that include a dimensionality reduction step (such as SCORPIUS, TSCAN, Monocle 2). Hence, we compared the performance of Monocle2 and TSCAN on their respective feature selection procedures and GO selected genes from redPATH. Results are shown in Figure S4 on four single cell datasets. The performance is significantly better in Figure S4 A-B when using GO selected genes in Monocle 2 and TSCAN, while the results are relatively similar on Dulken-NSC and Schlitz-HSC (Figure S4 C-D).

Additionally, we also compared redPATH against different gene selection methods (Figure S5). We evaluated redPATH 20 times on both GO selected genes and top 1,000 genes selected from dpFeature on six datasets. Although in most datasets, there is a drop in performance using dpFeature, the difference is mostly insignificant (as most achieved evaluation > 0.8). Interestingly, in mESC-Deng, redPATH managed to achieve better results with dpFeature (Figure S5D). In conclusion, we believe that our algorithm is relatively stable with different gene selection methods and will perform well in most datasets.

***Additional marker genes comparison between different algorithms***

*SOX9* and *APOE* are known NSC markers whilst *CCNA2* is a known marker for proliferation / cell cycle. Hence *CCNA2* can be used to distinguish between qNSC and aNSC. Identical to results shown in the main paper, SCORPIUS performs equally well in both NSC-Dulken and NSC-Llorens-B datasets compared to redPATH. However, the developmental trend in NSC-Shin differs. As shown in Figure S6A-B, qNSC is supposed to be highly expressed whilst SCORPIUS begin with relatively low expressed NSC at the beginning. Similarly, a contradicting trend is also depicted in *CCNA2*, violating the assumption of a linear development process from qNSC -> aNSC -> NPC -> NB.

***Heatmap analysis on MGH45 and MGH57***

Both MGH45 and MGH57 are WHO grade IV tumors. Within the malignant cells population, there is a clear small subpopulation in MGH57 while the expression change is relatively constant in MGH45 (Figure S9). The subpopulation inhibits some marker genes in glial cell differentiation / neuron fate commitment such as *OLIG1, OLIG2, IGFBPL1*.

***Modified Arbitrary Insertion Algorithm***

Given a graph **G**(V, E), where V is the vertices (or single cells in this case) and edges (KL distance between cells). A modification on the original algorithm [55] is given below:

1. Start with any arbitrary vertex $v_{i}$
2. Find vertex $v_{k}$ s.t. ${d(v}_{i}{,v}_{k})$ is minimized forming a tour $v_{i}- v_{k}- v_{j}$

*Selection Step:*

1. Select any arbitrary vertex $v_{k}$ currently not in sub-path

*Insertion Step:*

1. Find edge (*i, j*) in sub-path which minimizes

$${d(v}_{i}{,v}_{k})+ {d(v_{k}, v}_{j})- {d(v}_{i}{,v}_{j})$$

2. Insert $v_{k}$ between $v_{i}$ and $v_{j}$

Repeat selection step and insertion step iteratively until the Hamiltonian path solution is calculated.

**Supplementary Figures**

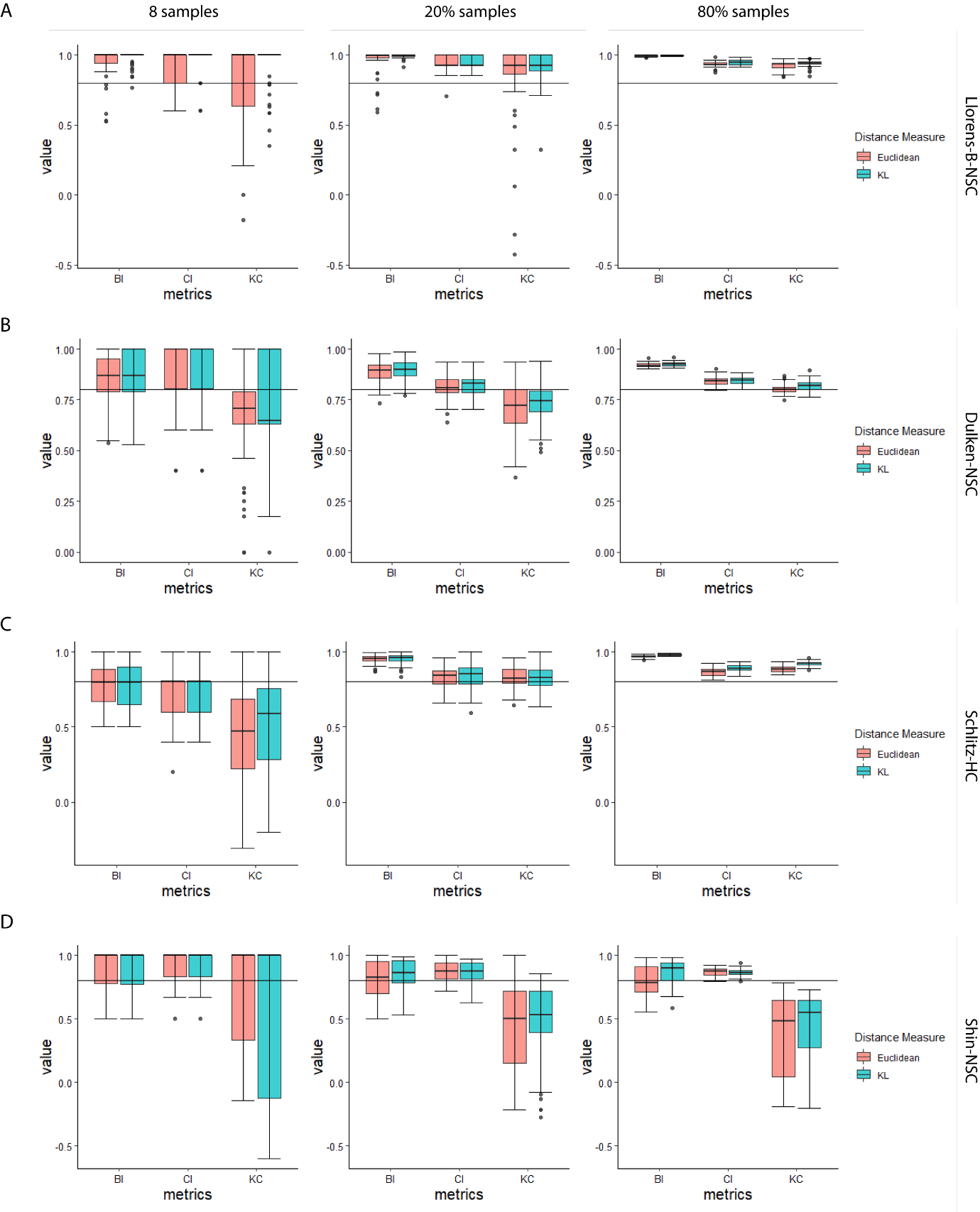

**Figure S1	Comparison of distance measures’ effect on redPATH**

**A-D)** Evaluation results of bubble sort index (BI), change index (CI), and Kendall correlation (KC) are shown for Llorens-B-NSC, Dulken-NSC, Schlitz-HC, and Shin-NSC respectively, comparing the difference in results between Euclidean distance and KL distance. Fifty different samples are taken for 8 cells, 20% of cells, and 80% of cells respectively for each column. A brute force Hamiltonian path is performed on 8 sampled cells, and redPATH is performed on the remaining two cases. Boxplot represents the quantile values based on the results from 50 random samples.

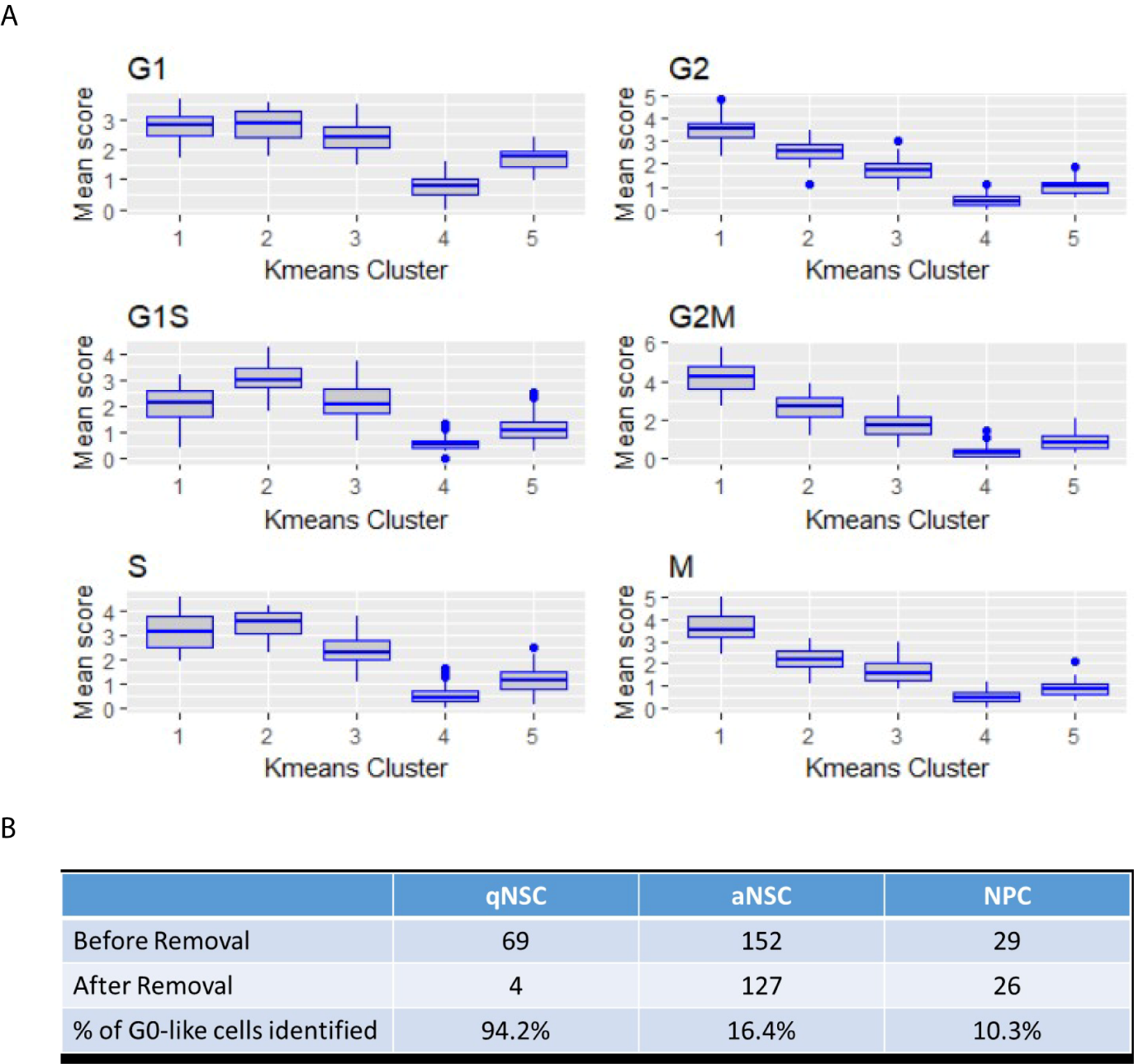

**Figure S2	Distribution of cycling scores and number of identified G0 cells**

**A** depicts the Dulken-NSC dataset showing the distribution of cell clusters in each of the six-cell cycle scores, namely G1, G2, S, M, G1S, and G2M. The lowest cluster is identified, and ANOVA test is performed to confirm that it is significantly lower than all other clusters in all six scores. **B** shows the percentage of G0-like cells removed in Dulken-NSC compared to the ground truth. qNSC is considered to be G0-like.

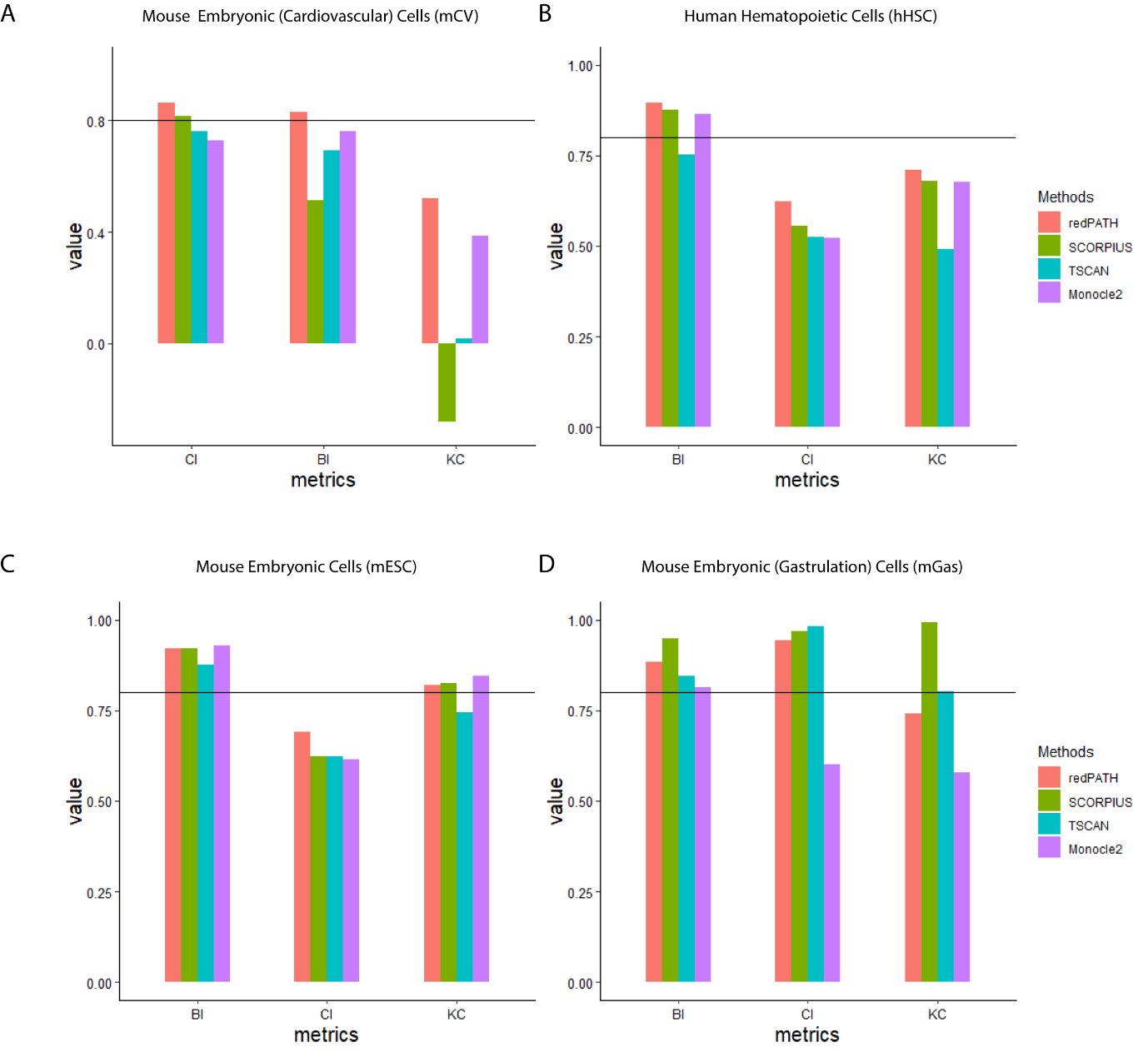

**Figure S3	Quantitative evaluation on multiple time point data**

**A-D)** Performance of each algorithm is evaluated on mCV, hHSC, mESC, and mGas, respectively. The horizontal line marks the 0.8 threshold for the evaluation metrics of BI, CI and KC. For each metric, bar plots representing the performance of each algorithm are plotted together for easy comparison.

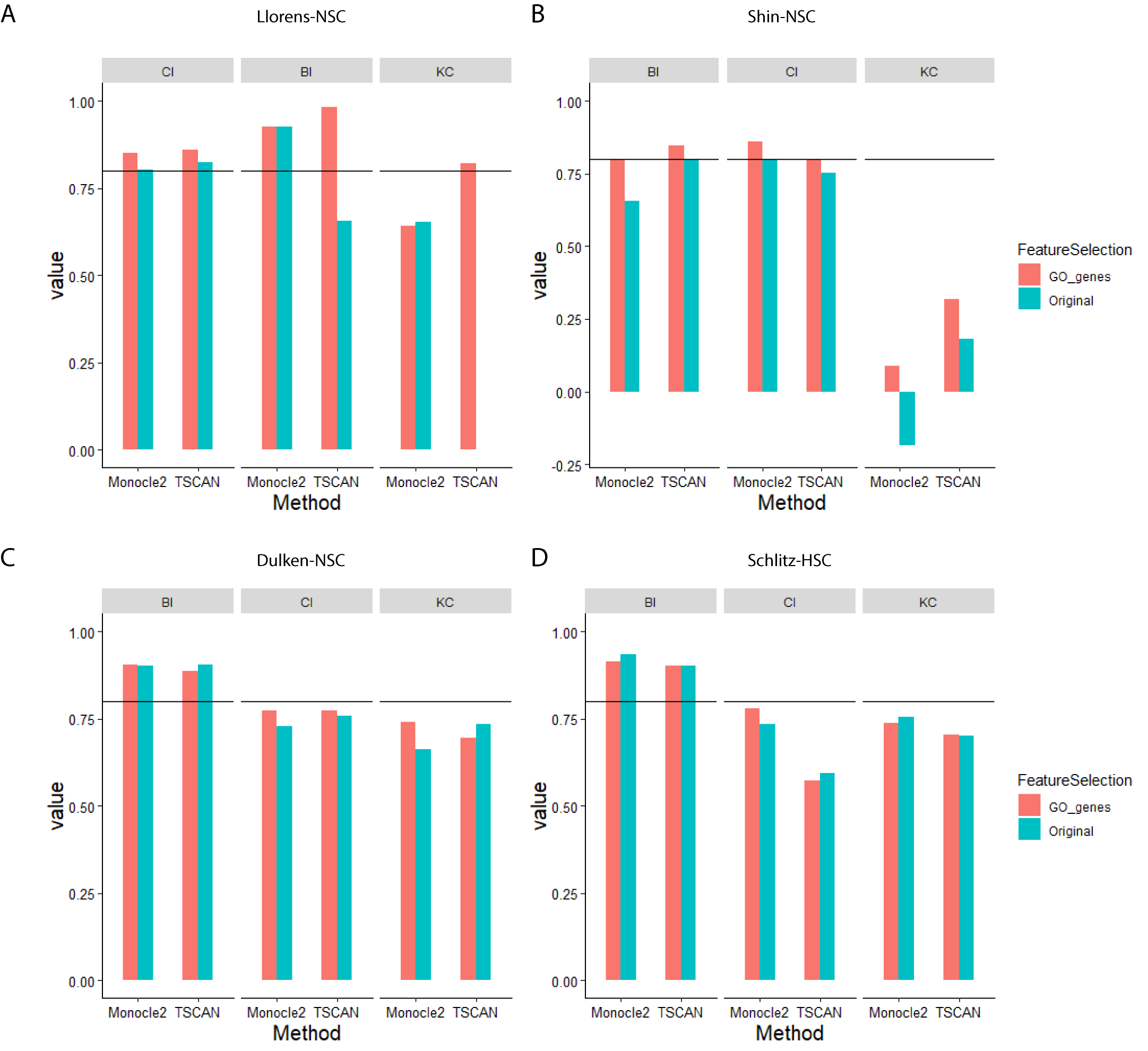

**Figure S4	Performance of different feature selection methods**

**A-D)** Performance of Monocle2 and TSCAN are compared using different feature selection methods on four validation datasets. Monocle2 uses dpFeature, and TSCAN uses its original pipeline for gene selection; both methods are compared against GO gene selection (from redPATH).

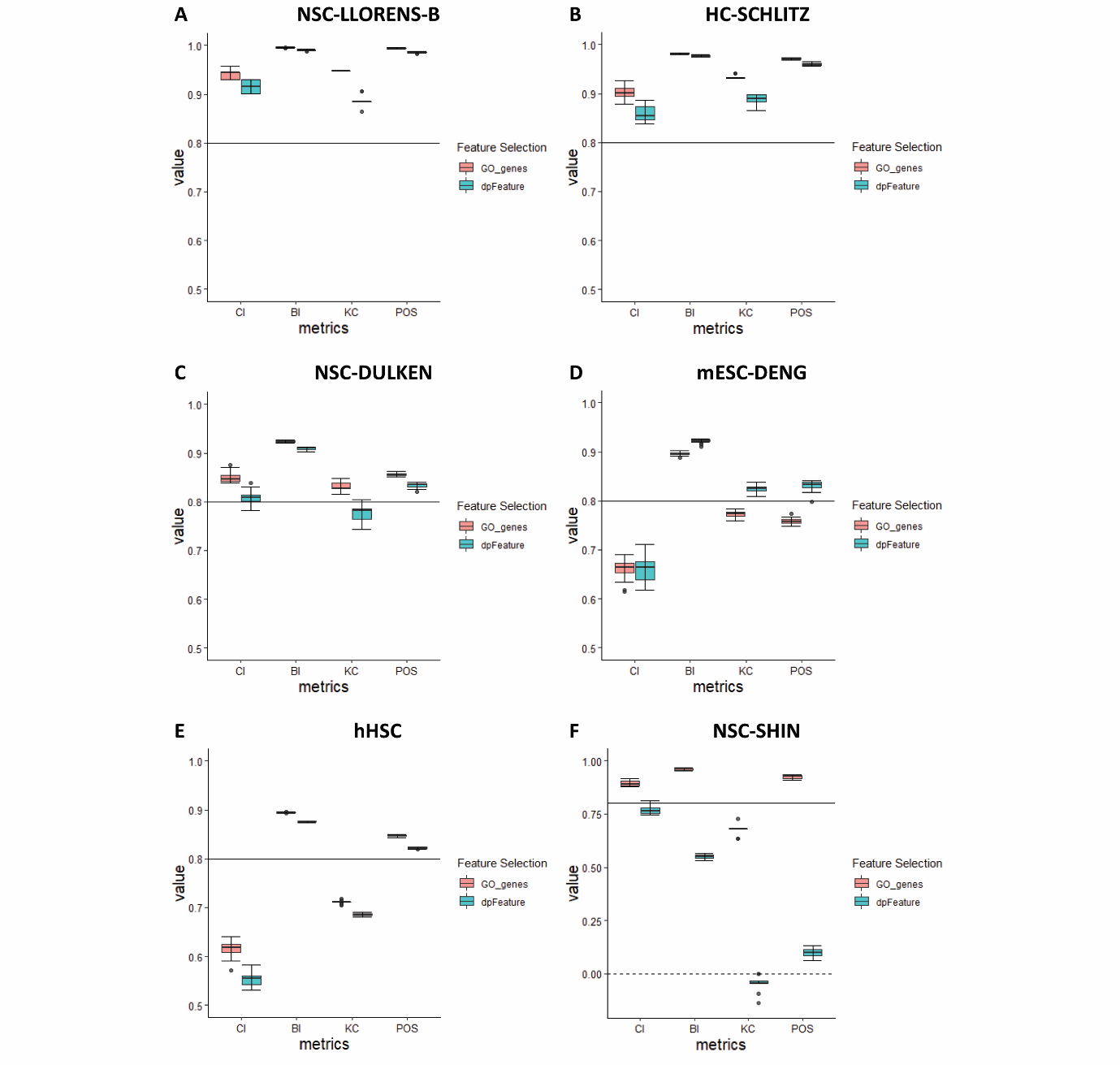

**Figure S5	Comparisons of feature selection on redPATH**

**A-F)** redPATH was performed 20 times for each feature selection method, namely GO genes (redPATH) and dpFeature (Monocle2). The performance was evaluated on four different evaluation metrics, change index (CI), bubble sort index (BI), Kendall correlation (KC) and POS score (POS). The solid horizontal line represents the 0.8 threshold value for evaluation, and the dotted horizontal line is set at 0. The dataset NSC-SHIN is plotted for range [-0.15, 1] while all the others were plotted in the range of [0.5, 1].

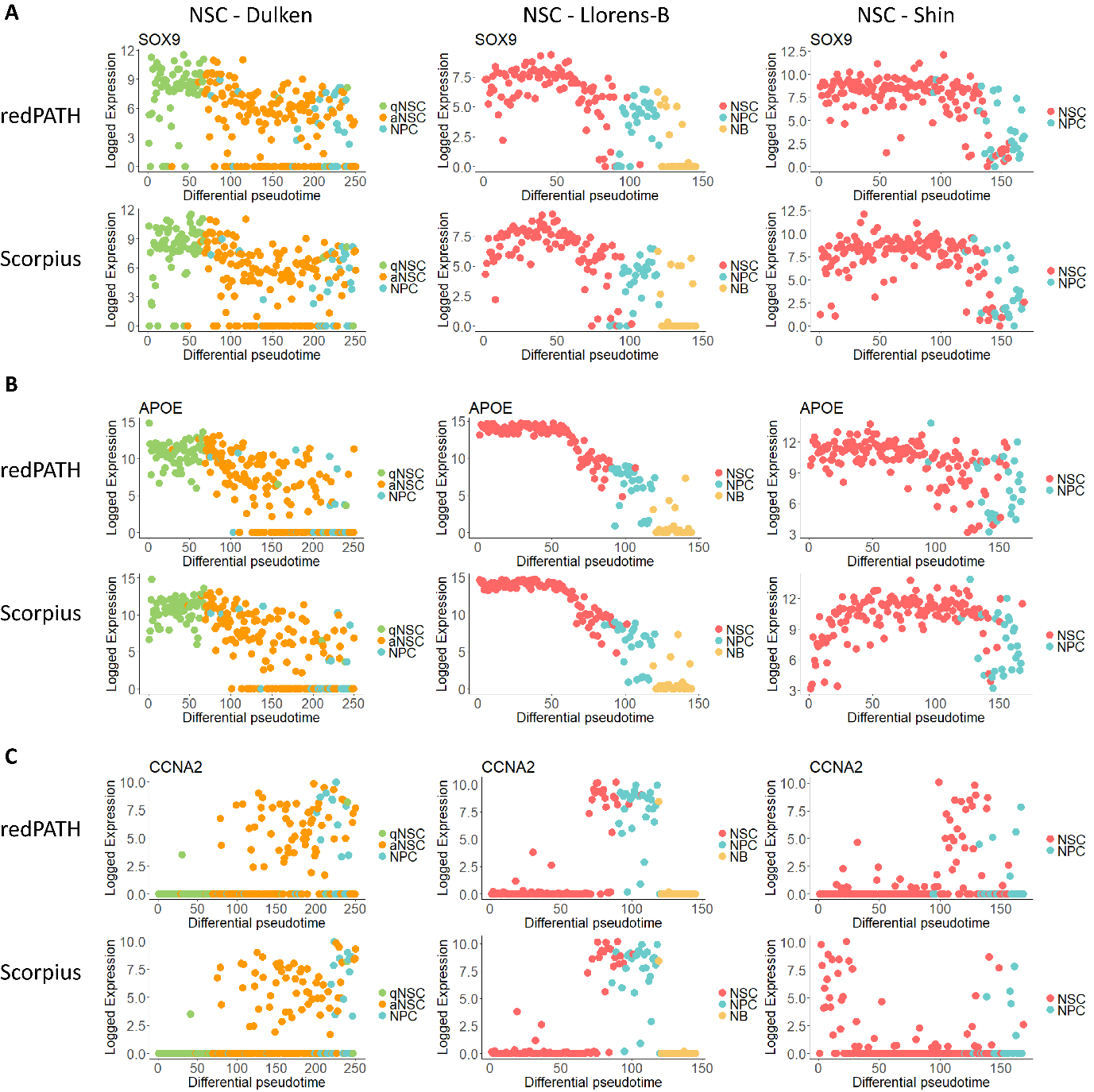

**Figure S6	Additional comparisons on NSC dataset**

The subtle differences in the inferred pseudo time are further compared on NSC marker genes (*Sox9, Apoe*) and proliferative marker gene (*Ccna2*). **A** depicts the difference in gene expression trend for *Sox9* by plotting the gene expression against inferred pseudo time. Comparison is made across three NSC datasets (Dulken, Llorens-B, and Shin respectively for each column) using redPATH and SCORPIUS. **B-C** Similarly for *Apoe* and *Ccna2,* respectively.

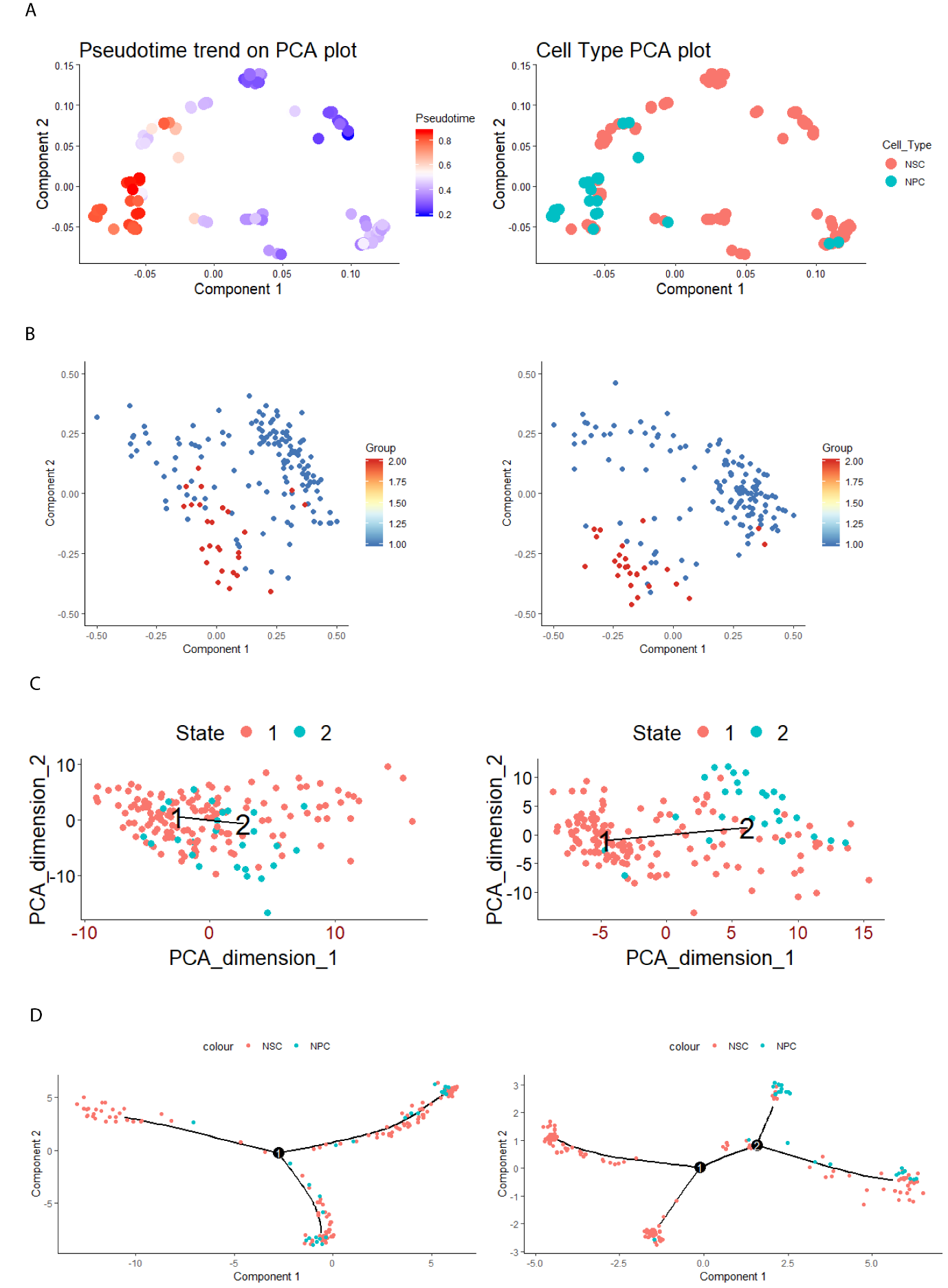

**Figure S7	Trajectory plots for Dulken-NSC**

Trajectory plots are generated for redPATH, SCORPIUS, TSCAN, and Monocle 2, respectively, using the default functions in their respective R packages. **A)** The left panel shows the pseudotime progression of cells, while the right denotes the cell type information. **B-D)** The left panel shows the trajectory obtained from using the default gene selection method in the respective package. On the right, the input expression is processed by using the GO selected genes from redPATH.

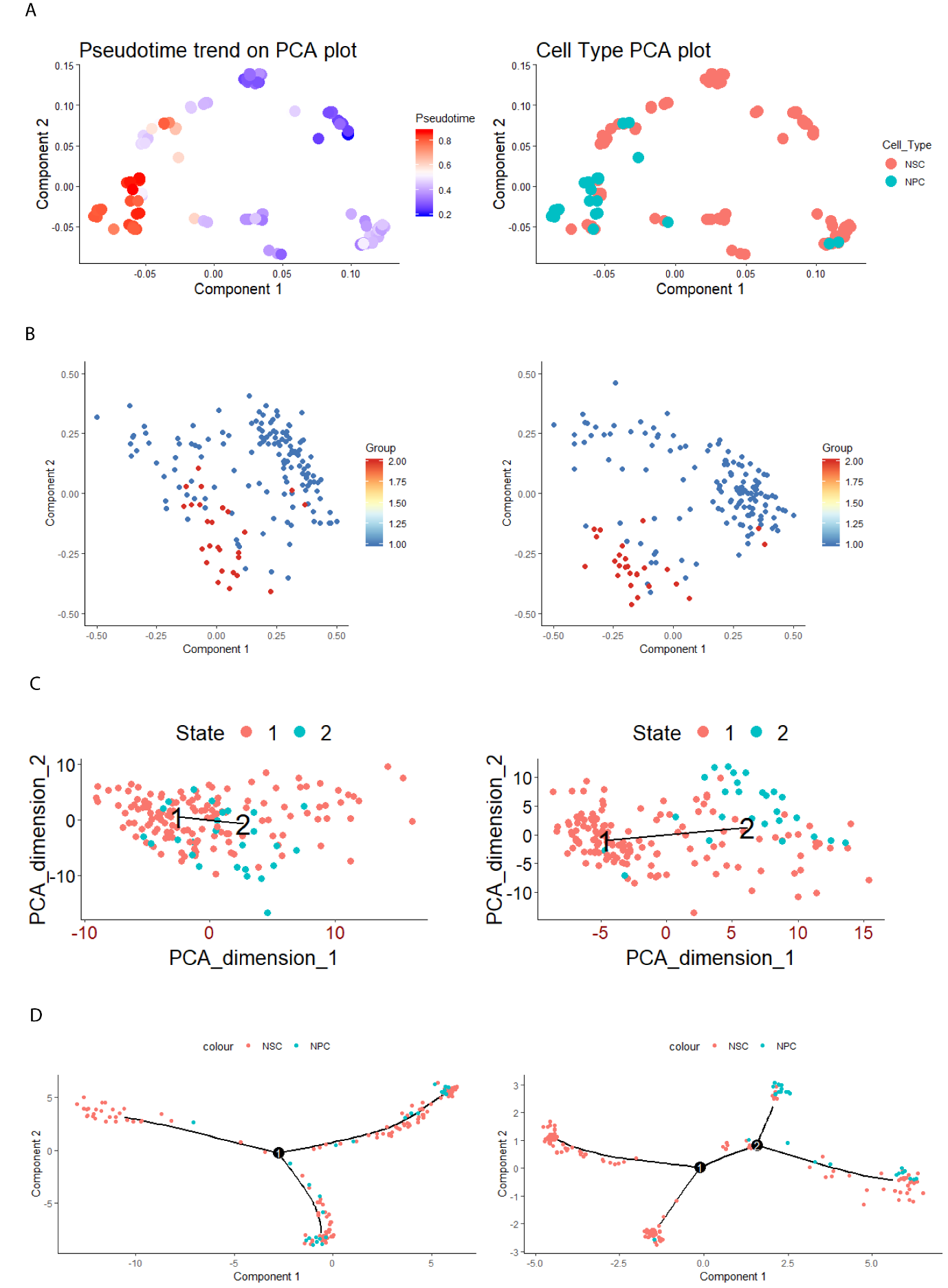

**Figure S8	Trajectory plots for Shin-NSC**

Trajectory plots are generated for redPATH, SCORPIUS, TSCAN, and Monocle 2, respectively, using the default functions in their respective R packages. **A)** The left panel shows the pseudo time progression of cells, while the right denotes the cell type information. **B-D)** The left panel shows the trajectory obtained from using the default gene selection method in the respective package. On the right, the input expression is processed by using the GO selected genes from redPATH.

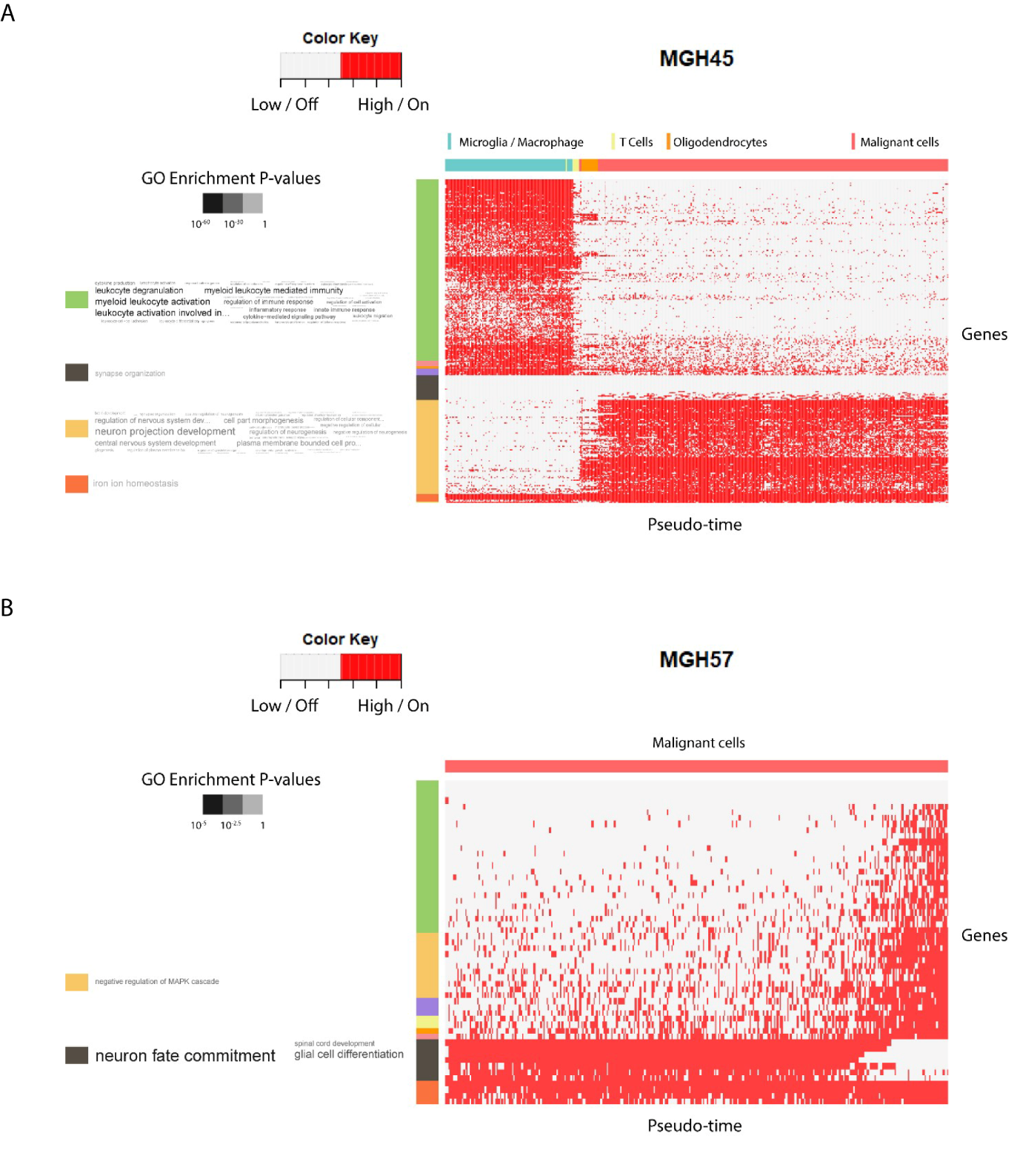

**Figure S9	Heatmap analysis on glioma datasets**

**A-B)** Heatmap plots for MGH45 and MGH57 respectively are produced according to the inferred HMM results from redPATH, indicating on / highly expressed state or off / lowly expressed state of each gene. The horizontal ordering denotes the differential pseudo time while each row represents a significantly identified gene. Gene clustering is shown on the left with Gene Ontology enrichments.

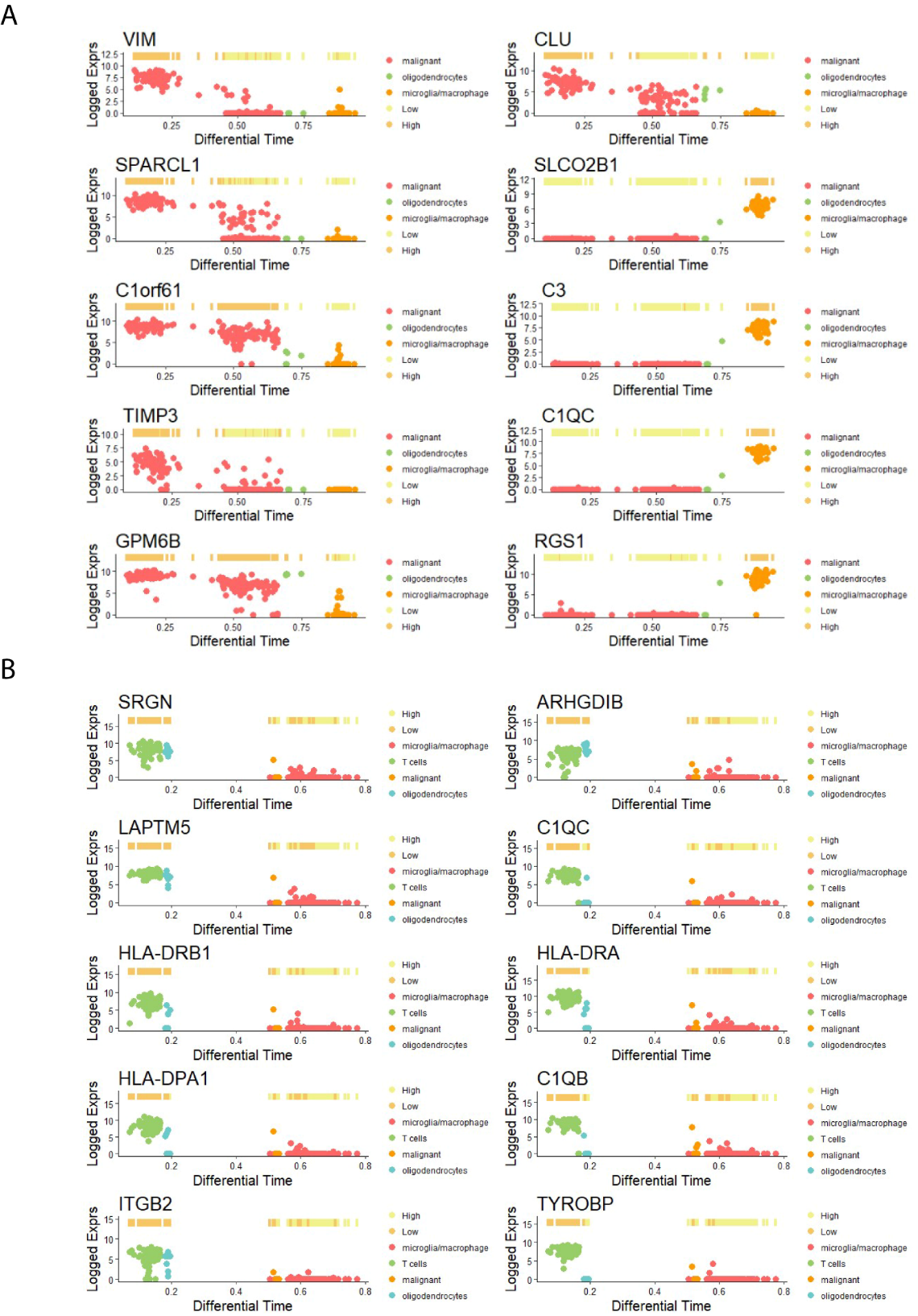

**Figure S10	Identified top rank genes from glioma datasets**

**A-B)** The top 10 genes are selected using dCor and MIC for MGH107 and MGH45, respectively. Each gene expression is plotted against the calculated pseudo time as well as the inferred state of the gene along the pseudo time. The cell type information is also reflected in the coloring of the points.
